# MethylNet: An Automated and Modular Deep Learning Approach for DNA Methylation Analysis

**DOI:** 10.1101/692665

**Authors:** Joshua J. Levy, Alexander J. Titus, Curtis L. Petersen, Youdinghuan Chen, Lucas A. Salas, Brock C. Christensen

## Abstract

**Background:** DNA methylation (DNAm) is an epigenetic regulator of gene expression programs that can be altered by environmental exposures, aging, and in pathogenesis. Traditional analyses that associate DNAm alterations with phenotypes suffer from multiple hypothesis testing and multi-collinearity due to the high-dimensional, continuous, interacting and non-linear nature of the data. Deep learning analyses have shown much promise to study disease heterogeneity. DNAm deep learning approaches have not yet been formalized into user-friendly frameworks for execution, training, and interpreting models. Here, we describe MethylNet, a DNAm deep learning method that can construct embeddings, make predictions, generate new data, and uncover unknown heterogeneity with minimal user supervision.

**Results:** The results of our experiments indicate that MethylNet can study cellular differences, grasp higher order information of cancer sub-types, estimate age and capture factors associated with smoking in concordance with known differences.

**Conclusion:** The ability of MethylNet to capture nonlinear interactions presents an opportunity for further study of unknown disease, cellular heterogeneity and aging processes.

## Background

Deep learning has emerged as a widely applicable modeling technique for a broad range of applications through the use of artificial neural networks (ANN) [1]. Recently, the accessibility of large datasets, graphics processing units (GPUs) and unsupervised generative techniques have made these approaches more accurate, tractable, and relevant for the analysis of molecular data [2–7].

DNA methylation (DNAm) is the addition of a methyl group to a nucleotide, typically cytosine, that does not alter the DNA sequence and occurs most frequently to cytosine-guanine dinucleotides (CpG). Methylated regions of DNA (hypermethylated), are associated with condensed chromatin, and when present near gene promoters, repression of transcription. Unmethylated regions of DNA (hypomethylated), are associated with open chromatin states and permissive to gene transcription. DNAm patterns are associated with cell-type-specific gene expression programs, and alterations to DNAm have been associated with aging and environmental exposures [8, 9]. Further, it is well-established that DNAm alterations contribute to development and progression of cancer. The hypermethylation of tumor suppressing genes and the hypomethylation of oncogenes can lead to pathogenesis and poor prognosis. Affordable array-based genome-scale approaches to measure DNAm have potentiated Epigenome Wide Association Studies (EWAS) for testing associations of DNAm with phenotypes, exposures, and states of human health and disease. Because DNAm patterns are cell-type specific, EWAS often account for potential confounding from variation in biospecimen cell composition using reference-based, or reference-free approaches to infer cell type proportions [10–13].

Measuring genome-wide DNAm in large numbers of specimens typically uses microarray-based technologies such as the Illumina HumanMethylation450 (450K) and HumanMethylationEPIC (850K) [14] arrays, which yield an approximation to the proportion of DNA copies that are methylated at each specific cytosine locus, and are reported as beta values. Preprocessing pipelines such as *PyMethylProcess* have simplified derivation and storage of methylation beta values in accessible data formats [15]. The scope of features from DNAm arrays is 20-50-fold higher than that of RNA-sequencing data sets that return normalized read counts for each gene. Though DNAm data can have a similar scope of features as genotyping array data sets, DNAm beta values are continuous (0-1), not categorical. Together, these facets of DNAm data sets pose challenges to analyses such as handling multi-collinearity and correcting for multiple hypothesis testing. To address these challenges, many downstream EWAS analyses have focused on reducing the dimensions into a rich feature set to associate with outcomes. By limiting the number of features through dimensionality reduction and feature selection, analyses become more computationally tractable and the burden of correcting for multiple comparisons is reduced.

An important advancement to methylation-based deep learning analyses was the application of Variational Auto-encoders (VAE). Initial deep learning approaches for DNAm data focused on estimating methylation status and imputation, performing classification and regression tasks, and performing embeddings of CpG methylation states to extract biologically meaningful lower-dimensional features [16–23]. VAEs embed the methylation profiles in a way that represents the original data with high fidelity while revealing nuances [4, 5, 24]. Thereafter, researchers attempted to develop similar frameworks for extracting features for downstream prediction tasks and identify meaningful relationships revealed by VAE latent representations [25]. However, VAE models are sensitive to the selection of hyperparameters [26] and have not been optimized for synthetic data generation, latent space exploration, and prediction tasks. Many auto-encoder approaches represent the data using an encoder, and then utilize a non-neural network model (e.g. support vector machine) to finalize the predictions. Presently, to the best of our knowledge there is no end-to-end training approach that both extracts biologically meaningful features through latent encoding and performs predictions using the derived features. Further, existing frameworks do not output predictions for multi-target regression tasks, such as cell-type deconvolution and subject age prediction.

Here, we leverage deep learning latent space regression and classification tasks through the development of a modular framework that is highly accessible to epigenetic researchers. *MethylNet* is a modular user-friendly deep learning framework for EWAS tasks with automation that leverages preprocessing pipelines. To discover important CpGs for each prediction we use the SHAP (SHapley Additive ExPlanation) approach [27]. We highlight *MethylNet* as an easy-to-use command line interface that utilizes automation to scale, optimize, and simplify deep learning methylation tasks. *MethylNet*’s capabilities are showcased here with unsupervised generative and clustering tasks, cell-type deconvolution, pan-cancer subtype classification, age regression, and smoking status classification. These analyses will pave the path for more robust deep learning prediction models for methylation data. Coupled with *PyMethylProcess* [15], we expect the *MethylNet* framework to enable rapid production-scale research and development in the deep learning epigenetic space.

## Results

Our approach uses a few simple commands, all of which can be executed for any prediction task. First, deep learning prediction models are pre-trained using variational auto-encoders, and the layers of the encoder are used to extract biologically meaningful features. These neural network layers are used to embed the data and extract features for clustering in the unsupervised setting, generating new data with high fidelity to the original source, and for prediction model pretraining. Second, prediction layers are included downstream of the encoder which fine-tune the model’s prediction and feature extraction layers end-to-end for the tasks of multi-output regression and classification. Training prediction layers optimize the neural network for prediction tasks. Third, autonomous hyperparameter scans are performed to optimize the model parameters for the first and second tasks while generating rich visualizations of the data. Lastly, the contribution of the CpGs to each prediction on varying degrees of granularity are determined through Shapley Feature Attribution methods.

*MethylNet* is implemented as a UNIX/Linux command-line tool that allows users to make deep-learning predictions on methylation data with use cases such as embedding, generation, classification and regression. With the specification of a single command-line option, *MethylNet* can be toggled between regression and classification tasks to address a wide breadth of problems. The modular, accessible characteristic of the MethylNet framework enables a simple procedure to train and produce results across multiple domains. In addition to predictive tasks, *MethylNet* can encode data into lower-dimension space from which to perform unsupervised clustering when researchers do not have labeled DNAm data. Further, MethylNet can generate realistic synthetic data with high fidelity relative to the original samples.

We show that *MethylNet* serves as an effective encoder for DNAm data by capturing latent features that have high fidelity to the original dataset. This method can utilize encodings to make accurate predictions in common DNAm analysis tasks, and the CpGs important for making predictions are concordant with prior observations. Finally, we demonstrate that *MethylNet* can also identify CpGs consistent with a large EWAS meta-analysis.

### Datasets Acquired

We selected three public DNAm data sets and use cases to illustrate a range of tasks and demonstrate ability to capture features that meaningfully encode aging, cell lineage, disease states, and exposures. The first dataset (Johansson data) was used to study both age and cell type classification and is one of the largest readily available DNAm datasets from healthy subjects with a wide age range (blood DNAm from individuals aged 15 to 95, GSE87571 [28]; Supplementary Figure 1 and Supplementary Table 1). The second dataset (The Cancer Genome Atlas, TCGA) was used to study cancer subtypes and includes 8,376 samples representing 32 different cancer subtypes (Supplementary Tables 1,2). The final dataset (Liu dataset) was used to compare blood DNAm in current smokers to never smokers among the controls from a rheumatoid arthritis study (GSE42861, subset n=188 [29]). All datasets were preprocessed using *PyMethylProcess* to yield 300k, 200k, and 300k CpG features respectively and then split into 70% training, 20% testing, and 10% validation.

### Evaluation of Unsupervised Encoder Performance

To establish *MethylNet* as a method for DNAm encoding, we first show that it can accurately capture features that dictate DNAm by using the features to recapitulate the supplied DNAm signal. If *MethylNet* can reconstruct the entire methylation profile from its latent-derived features, then those features should have high fidelity to the original dataset.

To support this, we first split up all of the CpGs into groups of features, with an average group size of 14,264 CpGs, that had shared methylation profiles across the training healthy blood samples of the Johansson data (n=503). Then, we fit an autoencoder produced by *MethylNet* to each of the resulting DNAm arrays and proceeded to test the ability to generate synthetic methylation samples on held out test data (Supplementary Figure 2). *MethylNet* was able to recapitulate the beta values with a weighted R^2^-value of 92.6% and weighted mean absolute error of 1.9% methylation between the original and generated held out test set (Supplementary Figures 2,3), demonstrating *MethylNet’s* ability to recapitulate the signal, and encode features with high fidelity to the original dataset.

To further establish encoding performance and provide confidence that future studies could utilize *MethylNet* to assess unknown heterogeneity in methylation profiles associated with disease, we tested whether features processed using the encoder can meaningfully cluster the methylation samples with concordance to known disease subtypes. TCGA DNAm data were encoded using *MethylNet’s* VAE and hierarchical clustering was performed on the embeddings and then compared with clustering results from Recursively Partitioned Mixture Modeling (RPMM)[30] that used 20k CpGs with the highest variance across samples. Cluster labels assigned to the embedded samples demonstrated agreement with the original cancer labels with a score of 0.76, compared to a score of 0.47 using RPMM (Supplementary Figure 4), indicating the potential for VAEs to utilize encoded features discover disease heterogeneity when labels are not supplied.

Given MethylNet’s performance in the unsupervised domain and its ability to meaningfully encode DNAm features, we next used this framework to validate performance in typical DNAm prediction tasks of age estimation, cellular proportion estimation, and disease classification.

### Age Results

DNAm-based age estimators such as the Horvath and Hannum clocks used elastic net penalized regression to identify sets of CpGs (353 and 71 respectively) strongly associated with age[31, 32]. Hannum et al. leveraged DNA methylation data from whole blood measured with the 450K Illumina platform in 656 subjects aged 19-101. Horvath leveraged genome-scale methylation data from 51 tissue and cell types in 82 independent data sets and over 8000 samples. The resulting models provide for very accurate age estimation but the number of and manner with which features can be associated with age are limited. Moreover, recently there is interest in understanding what drives observed remaining residual between chronological age and methylation age. The difference between age and methylation age has been termed biological age or age acceleration and has itself been associated with disease risk and all-causes mortality[33–35]. Demonstrating consistent performance between MethylNet and established approaches motivates future use of our method to study complex states and interactions underlying aging processes.

Again, utilizing the Johansson data, we trained MethylNet on the chronological age of the individuals to predict chronological age. *MethylNet*-predicted age showed excellent concordance with the actual subject age (R^2^=0.96, Figure 2a) in the hold-out test set (n=144), and only had 3.0 years mean absolute error (Figure 2b) (training and validation performance in Supplementary Table 3). These results are comparably accurate to those estimated by the Hannum and Horvath clocks. The contribution of each CpG to age groups binned by 10-year increments from ages 14 to 94 were measured by Shapley values. The CpGs with the one thousand largest Shapley values for each age group were overlapped with the CpGs of the Hannum clock (Figure 2c). These CpG contributions were compared between age groups using correlation distance, as illustrated in Figure 2d. The connectivity between different age groups’ CpG attributions in Figure 2d using hierarchical clustering demonstrates the sharing of important CpGs by similarly aged groups. Further description of the derivation of the Shapley score estimates can be found in the supplementary materials.

**Figure 1:**
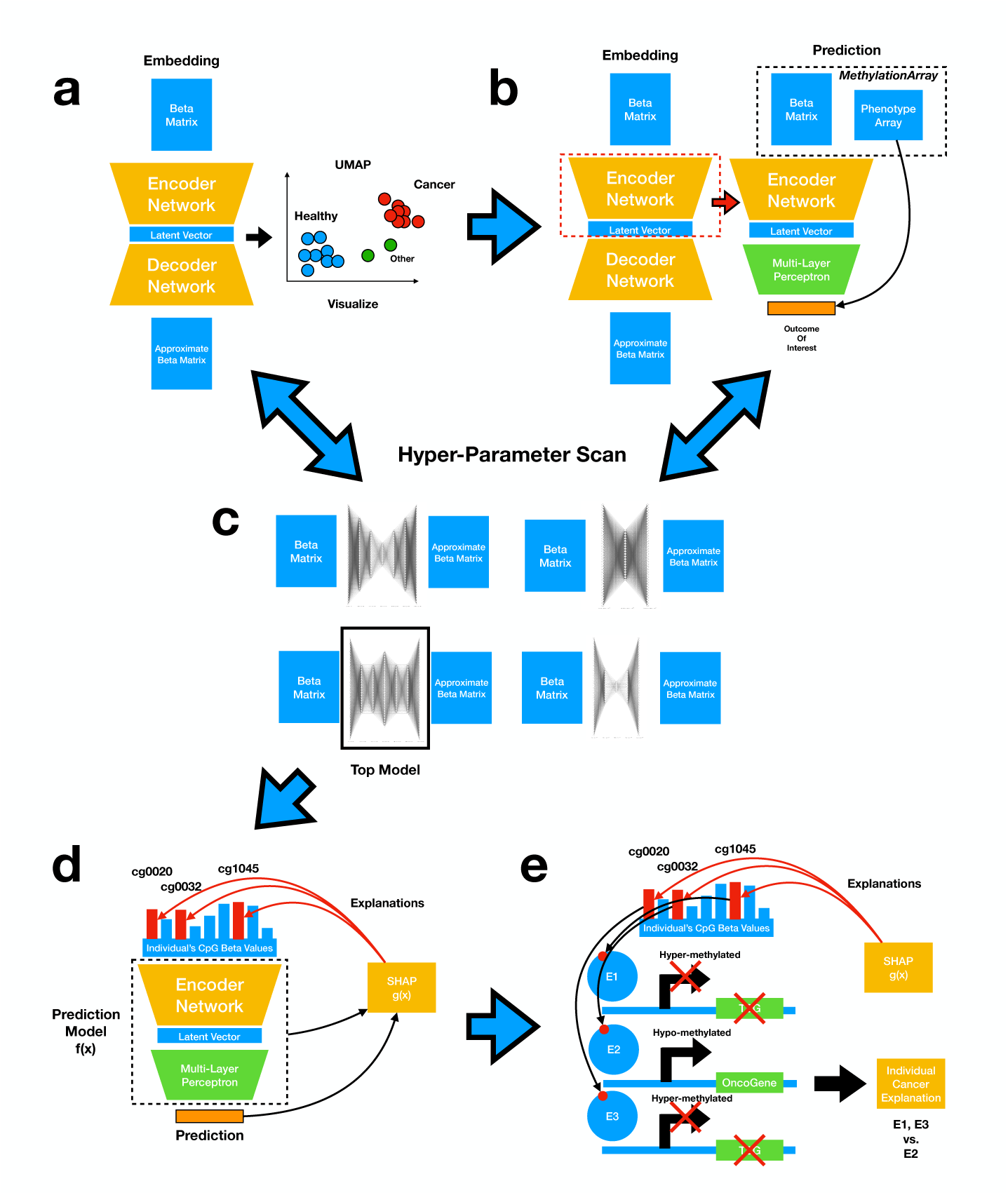
Step-by-step description of the modular framework: a) Train feature extraction network using variational auto-encoders; b) Fine-tune encoder for prediction tasks; c) Perform hyperparameter scans for (a) and (b); d) Identify contributing CpGs; e) Interpret the CpGs.

**Figure 2:**
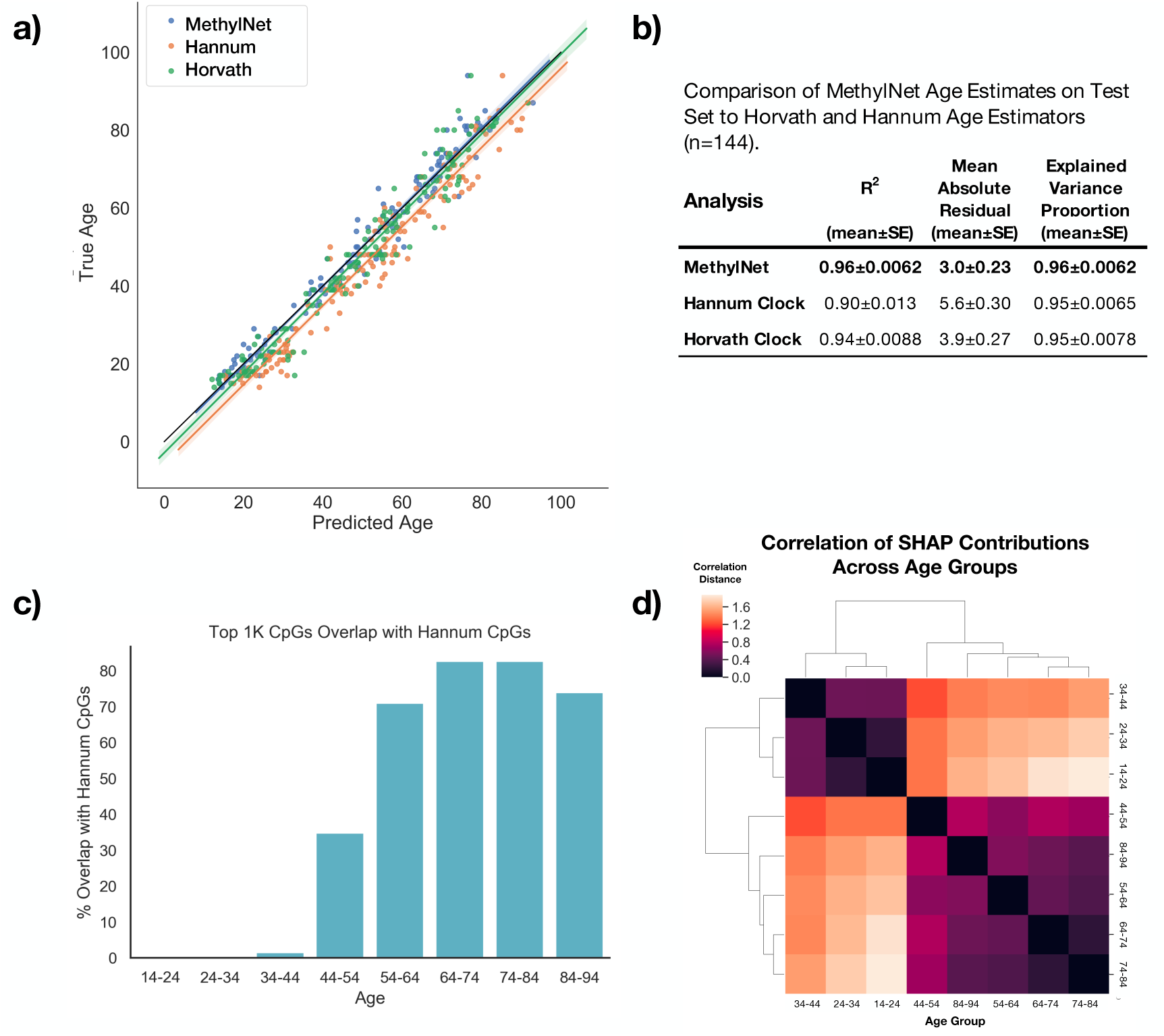
Age Results on Test Set (n=144): a) Age predictions derived using the Horvath, Hannum, and MethylNet estimators are compared to the true age of the individual, the predicted ages are plotted on the x-axis, the actual ages on the y-axis, and a line was fit to the data for each estimator; b) Comparison of MethylNet Age estimates on Test Set (n=144) to Horvath and Hannum Age Estimators. 95% confidence intervals for each score were calculated using a one thousand sample non-parametric bootstrap; c) Bar chart depicting the overlap of CpGs important to MethylNet and Hannum age estimators where one thousand CpGs with the highest SHAP scores per 10-year age group are divided by the total number of Hannum CpGs that passed QC; d) Hierarchical clustering using the correlation distance between SHAP CpG scores for age groups across all CpGs. The linkage is found between similar age groups.

We aimed to compare the highly contributing CpGs to age predictions using *MethylNet* and to those calibrated in the Hannum epigenetic clock [31]. The CpGs used by the Hannum model were most likely associated with those aged 60-80, the most prevalent ages in the cohort. Since the number of Hannum CpGs rediscovered by *MethylNet* appears to peak around this range, this supports evidence that *MethylNet* is able to recover the defining CpGs of the Hannum cohort.

### Cell Type Deconvolution Results

Reference-based cell type estimation approaches with DNAm data use a library of cell-specific leukocyte differentially methylated regions (L-DMR), to infer cellular proportions. These cell type libraries, similar to age estimation, contain a few hundred CpG features for prediction (e.g. the 350 CpG IDOL library[12]), and current deconvolution is very accurate and fast. Although current methods like estimateCellCounts2 accurately capture cellular proportions in blood, the future of cell type deconvolution includes efforts to estimate remaining sources of cell type heterogeneity, including cellular states that currently lack L-DMR. We sought to investigate the ability of MethylNet to capture current capabilities of cellular deconvolution so that it may be applied to future unsupervised domains when the requisite amount of data is available.

As such, *MethylNet* was tasked with estimating the cell-type proportions for six immune cell-types using the same dataset as supplied for the age analysis. As compared to the other EpiDISH estimator methods that utilize the IDOL library, the framework demonstrates exemplary performance on this task in R^2^ and mean absolute error across all cell-types save for monocytes, as demonstrated in Table 1 (Figure 3a-b; training and validation performance in Supplementary Table 4). Using Shapley attribution, contributions for each of the CpGs for driving the predictions of the cell-types was derived. Figure 3c shows the connectivity of their hierarchical clustering of these CpG attributions.

**Figure 3:**
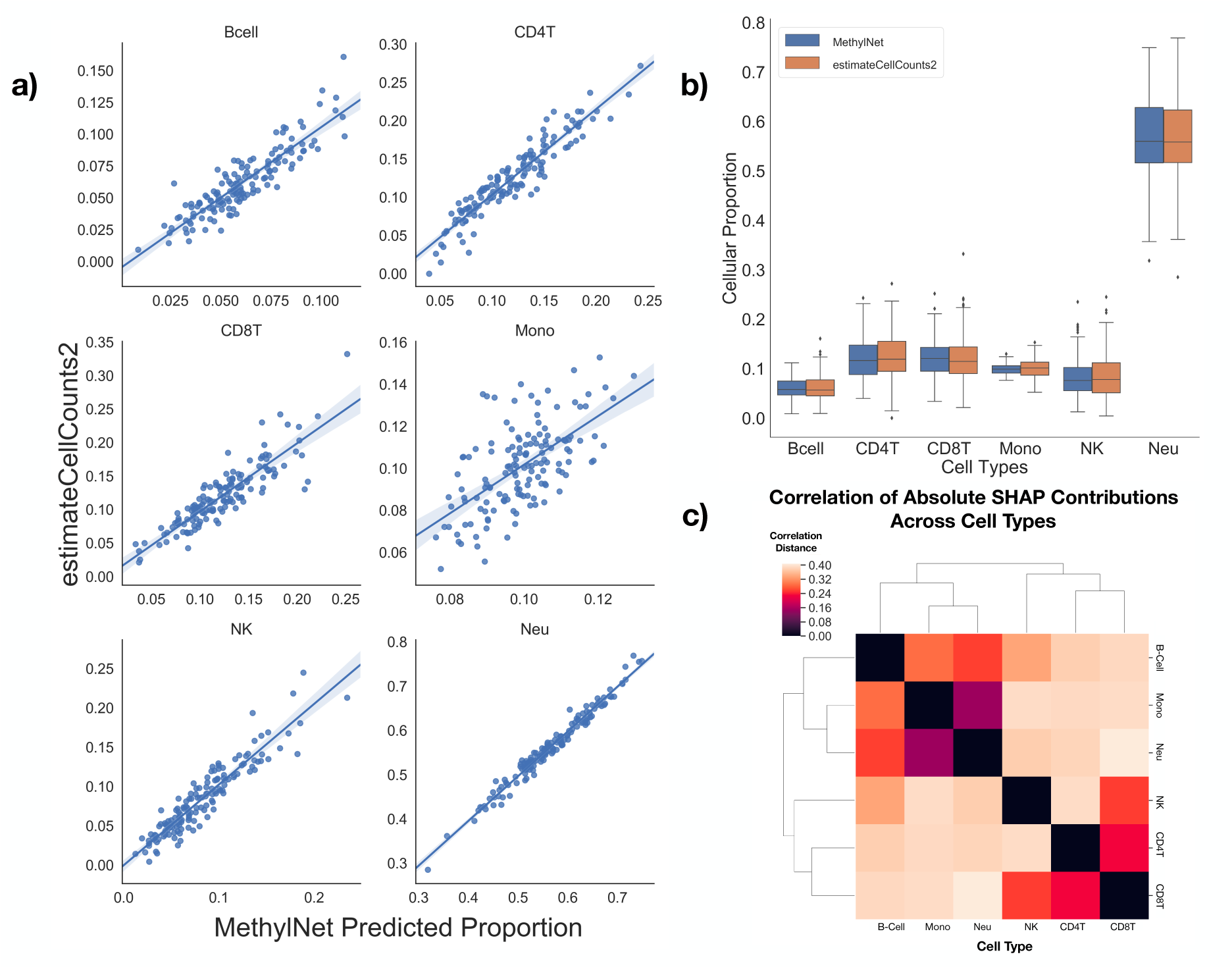
Results on test set (n=144) for cell-type deconvolution: a) For each cell type, the predicted cellular proportion using MethylNet (x-axis) was plotted against the predicted cellular proportion using estimateCellCounts2, which has been found to be a highly accurate measure of cellular proportions and thus serving as the ground truth for comparison, a regression line was fit to the data for each cell type: B-cell, CD4T, CD8T, Monocytes (Mono), NK cells, and Neutrophils (Neu); b) Grouped box plot demonstrating the concordance between the distributions of the MethylNet-estimated proportions of each cell-type and the distributions derived using estimateCellCounts2; c) Hierarchical clustering using the correlation distance between two cell types’ SHAP CpG scores across all CpGs. The linkage is found between cell types of similar lineage.

**Table 1.** Comparison of MethylNet Cell Type Deconvolution Results to IDOL Library EpiDISH Methods. 95% confidence intervals calculated using 1k-sample non-parametric bootstrap

| Library+Method | Cell Type | R<sup>2</sup> | Mean<br>Absolute<br>Residual | Explained<br>Variance<br>Proportion |
| --- | --- | --- | --- | --- |
|  |  | (mean±SE) | (mean±SE) | (mean±SE) |
| **300k+MethylNet** | CD8T | **0.78±0.04** | **0.016±0.0012** | **0.78±0.038** |
|  | CD4T | **0.86±0.018** | **0.014±9.0e-04** | 0.88±0.016 |
|  | NK | **0.87±0.017** | **0.012±8.5e-04** | **0.87±0.017** |
|  | B Cell | **0.79±0.026** | **0.009±6.3e-04** | 0.79±0.025 |
|  | Monocytes | 0.37±0.067 | 0.012±7.9e-04 | 0.38±0.062 |
|  | Neutrophil | **0.97±0.0043** | **0.011±7.1e-04** | **0.97±0.0042** |
| **IDOL+RPC** | CD8T | 0.72±0.061 | 0.019±0.0013 | 0.76±0.052 |
|  | CD4T | 0.34±0.091 | 0.036±0.0014 | **0.89±0.018** |
|  | NK | 0.024±0.11 | 0.033±0.0024 | 0.48±0.049 |
|  | B Cell | 0.77±0.035 | 0.01±5.3e-04 | **0.93±0.012** |
|  | Monocytes | 0.17±0.13 | 0.015±8.2e-04 | **0.64±0.053** |
|  | Neutrophil | 0.84±0.025 | 0.029±0.0013 | 0.96±0.0073 |
| **IDOL+Cibersort** | CD8T | 0.63±0.077 | 0.023±0.0014 | 0.75±0.055 |
|  | CD4T | 0.6±0.058 | 0.026±0.0014 | 0.86±0.02 |
|  | NK | -0.058±0.12 | 0.035±0.0025 | 0.46±0.055 |
|  | B Cell | 0.76±0.046 | 0.01±6.0e-04 | 0.88±0.024 |
|  | Monocytes | **0.45±0.089** | **0.012±7.2e-04** | 0.54±0.072 |
|  | Neutrophil | 0.91±0.015 | 0.02±0.0011 | 0.96±0.008 |

n=144, RPC: Robust Partial Correlations

The hierarchical clustering between the SHAP scores of each of the cell-types is consistent with the known cell lineage, reinforcing that cell lines that have co-evolved similarly share similar driving CpGs that are indicative of their cell-type. Some of the cell-types obtained improved concordance metrics (e.g. R^2^) compared to other cell types but had similar absolute errors (i.e. MAE). This is likely due to the fact that the total range of proportions of monocytes, for instance, from the collected data was small such that these errors could make it difficult to correlate the predicted and true cell type proportions. Alternatively, issues with the purity of the reference monocytes could complicate reference-library calibration. A similar overlap test was conducted between the *MethylNet* SHAP CpGs and IDOL-derived L-DMR CpGs (Supplementary Figure 5). Little overlap was found between the two sets, as only the B-cells were able to capture more than 10% of the IDOL CpGs. This does not indicate that *MethylNet* could not identify CpGs that are cell-type specific. Rather, this finding serves to indicate that models with different optimization objectives and number of features available differentially attribute CpGs.

To this point, we still do not know at what point do CpGs, across individuals or larger groupings reach statistical significance and thus warrant additional inspection. Some preliminary analysis can be found in the Supplementary Figures 7 and 8. For the Hannum and IDOL analysis, we set this at an arbitrary cutoff value of the top 1000 CpGs per age/cell-type group, but the distribution of these Shapley scores and their fidelity to model predictions is an active area of research [36].

### Pan-cancer Prediction Results

Finally, motivating uses of MethylNet as a mechanism to uncover sources of disease heterogeneity and the capability of the workflow to capture features that are tissue-specific, MethylNet was employed to make predictions of 32 cancer subtypes (n=1676) (one removed due to low sample size) across the pan-cancer TCGA cohort. This analysis yielded 0.97 accuracy, 0.97 precision, 0.97 recall and 0.97 F1-score, averaged across the different subtypes (Figure 4a) (training and validation performance in Supplementary Table 5). These results outperform a support vector machine (SVM)-based classification approach, in which *MethylNet* demonstrated a 0.15-unit (18%) increase in F1-score. A breakdown of classification accuracies for each subtype is in the supplemental results (Supplementary Tables 6,7).

**Figure 4:**
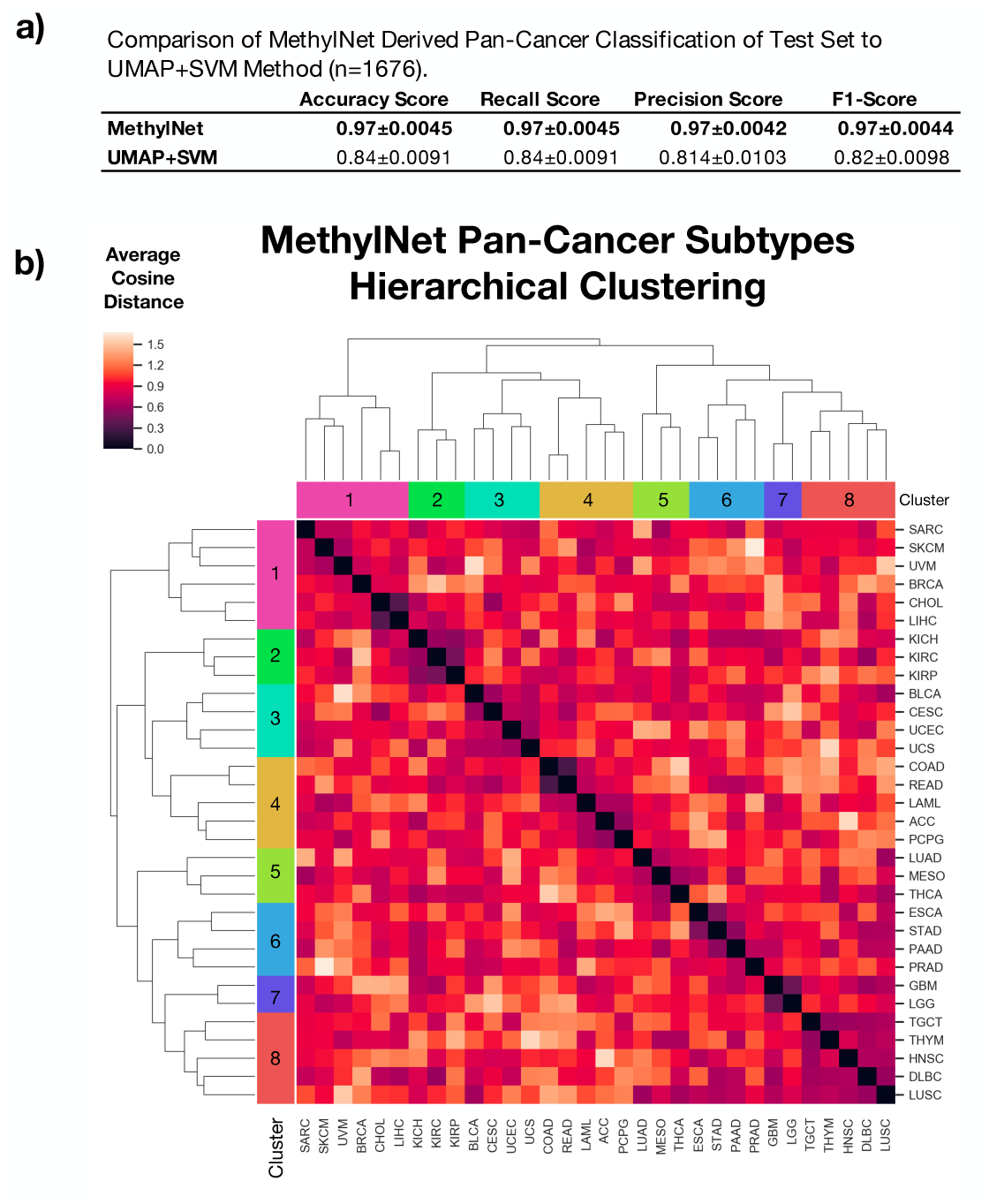
Results on test set for pan-cancer sub-type predictions: a) Comparison of MethylNet derived pan-cancer classification of test set (n=1676) to UMAP+SVM method. 95% confidence intervals for each score were calculated using a 1000 sample non-parametric bootstrap; b) Hierarchical clustering of average embedding cosine distance between all pairs of cancer subtypes. Cancer subtypes from both axes are colored by cancer superclasses, derived using the hierarchical clustering method. The clustering of similar MethylNet embeddings is concordant with known biology of tissue/cancer type difference. Skin and connective tissue cancers, and bile and liver cancers in Cluster 1. All kidney cancers in Cluster 2. Bladder, uterine and cervix cancers in Cluster 3. Pairing of colon and rectal cancers, both adrenal cancers in Cluster 4. A tie between lung adenocarcinoma and mesothelioma in Cluster 5, both of which may develop in similar locations. Pairings between stomach and esophagus cancer, and pancreas and prostate cancers in Cluster 6. Brain cancers in Cluster 7. Thymoma, Diffuse Large B-Cell lymphomas in Cluster 8. While the lung cancers were not paired together, they experienced a high degree of embedded similarity. The connectivity between the lung squamous cell cancer and its neighboring types prevented the two cancers from being grouped together.

The latent profiles derived for pan-cancer subtypes given the model training on this predictive task showed clustering with high concordance to known cancer type differences. Thresholding a hierarchical clustering of the average cosine distance between cancer subtypes from the *MethylNet* derived embeddings (Figure 4b, Supplementary Table 8) indicates clustering of the test methylation profiles by eight unsupervised biologically corresponding superclasses. The subtypes that define these larger groupings are concordant with expectations from tissue differences in cancer biology.

Taken together, *MethylNet* not only makes highly accurate and robust classification predictions, but also extracts latent features with high fidelity to the biology of tissue or cancer type difference.

The similarity between some of the subtypes may explain why and how certain subtypes did not perform as well compared to others (Supplementary Tables 6 and 8). For instance, we see that 4 KIRC and KIRP cases were conflated with each other. In addition, two cervix cases were predicted to be uterine. There were elevated rates of misclassification between the colon and rectal cancer pairings and esophageal, head and neck, and stomach cancer pairings. Finally, seven predicted glioblastoma cases were actually low-grade glioma (Supplementary Table 6). Thus, subtypes tended to be misclassified only within each superclass. The exception to this trend was the misclassification of lung squamous cell carcinomas, four of which were predicted to be its adenocarcinoma counterpart, which is consistent with the shared embedding profile, and likely reflects similar biology of cellular lineage.

For the cancer subtype analysis, we sought to identify concordance between the latent profiles of methylation across cancer types. Because each tumor type has a different baseline DNAm profile for its normal tissues-of-origin, and these differences are expected to contribute to the prediction, we decided not to attempt derivation of the salient CpGs for each subtype’s prediction.

### Dataset Scaling and Comparison to Multi-layer Perceptron

In addition to evaluating the disease subtypes, we sought to use the TCGA pan-cancer dataset to better elucidate the framework’s sensitivity to reductions in number of features and number of training samples. Performance was observed to increase linearly with number of training samples and logarithmically with available CpGs (Supplementary Figure 9). We also demonstrate comparable performance of training a multi-layer perceptron using similar training parameters and neural network architecture as *MethylNet’s* encoder-prediction structure (Supplementary Figure 9). *MethylNet’s* encoders have utility for generative and unsupervised tasks through capturing a low dimensional distribution of the data and the strengths in initializing the early layers of the prediction model with encoder pretraining. We have also included an implementation of the multi-layer perceptron that can be trained within our framework.

### EWAS Application

Given the success of *MethylNet* to capture nonlinear interacting features that cluster, recapitulate and assist with predictions, we sought to evaluate *MethylNet* on the Liu data for the prediction of smoking status (current vs. never smoker) and compare the results to a prior robust EWAS meta-analysis [37]. *MethylNet* achieved 73% accuracy in predicting smoking status despite relatively small training (n=139), validation (n=19) and held-out test sets (n=30) (Supplementary Figure 10). There was a significant correlation between the rank of CpGs most important in differentiating smoking status found through MethylNet (average SHAP ranking for each CpG) and the rank of the CpGs significantly associated with smoking using a significantly larger dataset by Joehanes et al. (r=0.69; p-value=0 for statistical test of non-correlation) (Supplementary Figure 10). The preservation of these ranks indicates that *MethylNet* can form associations with outcomes that are concordant to known EWAS analyses, even though it places more emphasis on interacting features versus the traditional EWAS.

## Discussion

Here, we introduce *MethylNet*, a modular deep learning framework that is easy to train, apply, and share. *MethylNet* employs an object-oriented application programming interface (API) and has built-in functionality to easily switch between analyses with respect to embedding, generation, classification, and regression tasks. We demonstrate *MethylNet’s* ability to capture features that recapitulated the original DNAm data and generated accurate predictions that conform with expected biology. *MethylNet* extends previous approaches by fine-tuning the feature extractor and adding additional layers for prediction tasks. It also employs a robust hyperparameter search method that optimizes the parameters of the model for generalization to unseen data. The pipeline is flexible to the demands of the user. For instance, if a user only wanted to train a custom machine learning model on the latent features, the data can be extracted before the end-to-end training step. By demonstrating the ability to meaningfully encode DNAm features, predictive performance on four tasks; age prediction, cell-type deconvolution, pan-cancer subtype prediction, and concordance to the results of a known EWAS meta-analysis; we present further support of the applicability of VAEs for feature extraction, and more evidence that deep learning presents an opportunity for learning meaningful biology and making accurate predictions from feature-rich molecular data.

### Strengths, Limitations, and Future Directions

Interpretation of our high dimensional models still has challenges, partially due to the drawbacks of assigning feature attributions to high dimensional multi-collinear data. While traditional linear models can still be highly predictive, multi-collinearity has the effect of adjusting the coefficients of the predictors such that the results are not as interpretable.

Shapley feature attributions are a promising method used to explain predictions estimating complex models with simpler linear ones as we able to demonstrate agreement between age groups and cell lineages and concordance between ranked SHAP scores and ranked p-values of CpGs associated with smoking status of a large EWAS meta-analysis.

Our age and cell-type analyses were conducted to demonstrate the capabilities of the deep learning tool and models were trained on a relatively small study of blood samples, only a subset of those included in the Horvath framework.

Further work can capture features indicative of age acceleration, a popularized prognostic indicator tied to the residual between the predicted and actual age. Since initial publications in 2013, investigators have started using the difference between chronologic and predicted DNAm age to investigate questions related to so called biological age or age acceleration[38]. This area of epigenetics is moving towards understanding the relation of the age residual with disease risk, and potential to modify it through intervention (e.g. diet and exercise). More advanced treatment of the data underlying prediction of age will allow opportunities for mechanistically informed intervention studies that aim to reduce age acceleration and improve public health [9].

*MethylNet* methodology presents alternative framework to uncover functional gene regulation that accounts for biological age acceleration and goes beyond the limited set of features used to predict methylation age in Horvath, Hannum, and other DNA methylation clocks. As the biology of these clocks are still being discovered [39] and due to the non-linear relationship with both chronological age [40] and other biomarkers of cell epigenetic cell maturation[41], further examination of age acceleration and biology should be done through neural networks.

Our analyses also only presented predictions across one type of tissue without yet accounting for differences in methylation between cell types. *MethylNet* was shown to capture some of the remaining sources of cellular heterogeneity, which can include differential methylation of cell subtypes and states that are known to exist, but for which we do not currently have L-DMRs.

*MethylNet* represents an opportunity to improve reference-based and reference-free deconvolution approaches. More robust and consistent estimators that address current limitations of DNAm-based deconvolution approaches will be the focus of future applications of the *MethylNet* method.

Prior works that have explored pan-cancer prediction in the deep learning space have limited their analyses to a small set of CpGs that do not capture a holistic understanding of interaction and regulation in the cancer context[42]. Our results demonstrate that models with a larger number of CpGs are needed to accurately capture differences in tissue/cancer subtypes. Since *MethylNet* captures and confounds the biology between similar conditions, it presents an opportunity to explore similar therapeutic targets and treatments across disease types of similar tissue, within and outside cancer studies. Given the ability of *MethylNet* to capture the differences in the profiles between the cancer subtypes, there is great opportunity to better understand heterogeneity of other diseases.

Our analyses refrain from uncovering relationship between the discovered CpGs and functional effects because of the difficulties associated with localizing the effect of a small set of CpGs of interest. Once the salient attributions are found, CpG analyses experience common pitfalls when trying to match CpGs to their nearest gene via the found promoter region. Such analyses may ascribe the CpG’s effect in the context of what gene they appear to be regulating.

However, genes are also regulated at a distance in the 3D topological space by interacting with enhancer regions [43, 44]. Thus, enrichment methods based on individual gene to CpGs relationships implemented in missMethyl[45] may not be suitable for interpreting loci identified by *MethylNet.* Ideally, downstream approaches to add biological interpretation would take into account chromosome/genome interaction (e.g. through use of Hi-C data) and genome topological structure/organization. For instance, enrichment from chromatin state and histone modifications present in the target loci as used by ChromHMM and LOLA [46, 47] might be more warranted. Some model result interpretation issues may be partially circumvented by integrating gene expression data into the model or more structurally by building a deep learning mechanism to predict gene expression from DNA methylation using other layers of information from the genomic context [48].

An important take-away is that as interpretation methods for these high dimensional data are pioneered, VAE-based deep learning models will likely find CpGs that interact in ways we would not traditionally think about. While the other models were trained on a much smaller set of CpGs, *MethylNet* is able to make its predictions on 200-300K CpGs, capturing complex interactions between a much larger set of CpGs. Crucial next steps should address these interpretability and confounding concerns through feature selection, covariate adjustment and more biologically interpretable informatics methods for CpG interpretation.

Finally, to scale up *MethylNet’s* deep learning workflows to production grade as well as incorporate information from Whole Genome and Reduced Representation Bisulfite Sequencing, future renditions may utilize common workflow language (CWL) [49]. In addition, new Bayesian search methods may be employed to better automate the selection of model hyperparameters and automate the construction of the ideal neural network architecture [50, 51].

## Conclusion

We demonstrate a modular, reproducible, and easy-to-use object-oriented deep learning framework for methylation data: *MethylNet*. We illustrate that *MethylNet* captures meaningful features that can be used for future unsupervised analyses and achieves high predictive accuracy across age estimation, cell-type deconvolution, cancer subtype, and smoking status prediction tasks. *MethylNet’s* accuracy at these tasks was superior, or at least equivalent to, other methods and interpretations of the model’s outputs demonstrated agreement with prior literature. We hope that *MethylNet* will be used by the greater biomedical community to rapidly generate and evaluate testable biological hypotheses involving DNA methylation data through a scalable, automated, intuitive, and user-friendly deep learning framework.

## Methods

### Description of Framework

Here, we present a description of a modular and highly accessible framework for deep learning tasks pertaining to unsupervised embedding, supervised classification and multi-output regression of DNA methylation (DNAm) data. The *MethylNet* pipeline comprises modules and commands specifically pertaining to embedding, prediction, and interpretation.

First, after preprocessing using *PyMethylProcess*. The dataset is split into training, validation, and testing sets using *train_test_val_split* of the preprocessing pipeline utilities.

### Training the Feature Extractor to Embed Data

The embedding module is used to pretrain the final prediction model by using Variational Autoencoders to find unsupervised latent representations of the data. Pre-training is an important part of transfer-learning applications. The knowledge extracted from learning unsupervised representation of the data is used towards learning predictive tasks with a lower data requirement. Data fed into these VAEs pass through an encoder network that serves to compress the data and then this compressed representation is fed into a decoder network that attempts to reconstruct the original dataset while attempting to generate synthetic samples. The model attempts to balance the ability to generate synthetic samples with the ability of the data to be accurately reconstructed. The weight given to generation versus reconstruction can be set as a hyperparameter [52]. Generating synthetic training examples are important for adding noise while training a network for prediction tasks, a component which serves as a form of regularization to make the algorithm more generalizable to real-world data. While synthetic data can be generated using *MethylNet* via the *generate_embed* command, this generative process is meaningfully utilized during training, when the algorithm samples from the latent distribution of the embedded data to regularize. Nevertheless, the ability to reconstruct the original dataset is important because it governs how latent representations of the data are capturing features that properly describe the underlying signal.

In order to run the embedding module on the input *MethylationArray* training and validation objects, *perform_embedding* is executed via the command line interface. Hyperparameters of the autoencoder model can be scanned via the *launch_hyperparameter_scan* command. This randomly searches a grid of hyper-parameters and randomly generates neural network topologies (number of layers, number of nodes per layer). The complexity (network width and depth), of which can be weighted by the user. The framework stores the results of each training run into logs to find the model with the lowest validation loss (Binary Cross Entropy reconstruction loss plus KL-Loss of the validation set) (hyperparameters with lowest validation loss can be found in Supplementary Table 9). Alternatively, results from the embedding module can be input into any machine learning algorithm of choice. Embedding results are visualized through interactive 3-D plots by running *transform_plot* from *PyMethylProcess*.

### Training for Prediction via Transfer Learning

*MethylNet* can be used to perform classification, regression, and multi-output regression tasks via the prediction module. The prediction module uses *MLPFinetuneVAE* to fine-tune encoding layers of VAE model while simultaneously training a few appended hidden layers for prediction. The *make_prediction* command is run for these prediction tasks, and hyper parameters such as model complexity and learning rate and schedulers are scanned via the *launch_hyperparameter_scan* module (hyperparameters with lowest validation loss can be found in Supplementary Table 10). The final model is chosen if it has the lowest validation loss (Mean Squared Error for Regression, Cross-Entropy for Prediction), and the output model is a snapshot at the epoch that demonstrated the lowest validation loss. The test set is also evaluated immediately after the model is trained using the training set. The results from *MethylNet* can be immediately benchmarked and compared for performance to other machine learning algorithms, which can be evaluated using the *general_machine_learning* module from *PyMethylProcess*. Furthermore, ROC Curves and classification resorts can be output using *plot_roc_curve* and *classification_report* and regression reports are generated via *regression_report.* A confusion matrix of misclassifications can be generated from *PyMethylProcess*’s *plot_heatmap*. Finally, the training curves for both the embedding and prediction modules can be visualized using the *plot_training_curve* command (example prediction embedding plots found in Supplementary Figure 6; analysis training curves can be found in Supplementary Figure 11).

### Interpretation of Results

Predictions from *MethylNet* can be interrogated in two ways. The first approach uses SHAPley feature attribution to assign a contribution score to each CpG based on how much it contributed to the prediction. The second approach compares learned clusters of embeddings of methylation samples (and corresponding subtypes), for biological plausibility.

The SHAPley value interpretations, available using *methylnet-interpret* approximate the more complex neural network model using a linear model for each individual prediction, the coefficients of which are Shapley values. Shapley values represent the contributions of each CpG to the individual predictions. They are produced after the prediction model and test *MethylationArray* are input to the *produce_shapley_data* command, which dumps a *ShapleyData* object into memory. The Shapley coefficients can be averaged by condition to yield summary measures of the importance of each CpG to the coarser category, and the coefficients can be clustered to demonstrate the similarity between methylation subtypes and coarser conditions, which can be compared to known biology.

### Description of Experiment

We evaluated our *MethylNet* framework (hyperparameter scan, embedding, fine-tuning predictions, interpretation) using 34 datasets from n=9,500 samples for four different prediction tasks: classification (TCGA pan-cancer subtype and smoking prediction), regression (age prediction), and multi-output regression (cell-type deconvolution).

*PyMethylProcess* was used to preprocess the data, and yielded *MethylationArray* objects that contain a matrix of beta values for each individual and the corresponding phenotype information [15]. The *MethylationArray* data for each of these three experiments were split into 70% training, 20% testing, and 10% validation sets. The training set was used to update the parameters of the model. The validation set was used to terminate training early and choose hyperparameters that would be most generalizable to a test set. The test set was used for final model evaluation and interpretation. More information on model training can be found in the supplementals. For each score, 95% confidence intervals were computed using a one thousand sample non-parametric bootstrap.

First, *MethylNet*’s generative analysis was conducted on 8 arrays representing 8 groupings of features of the Johansson data, found by running a KMeans clustering algorithm on a UMAP clustering of CpG Methylation profiles. Each group was trained using a 50-job VAE hyperparameter scans to yield the ideal embedding. A *generate_embed* command was used to first embed methylation profiles and then decode them to their predicted values. All of the beta values of the CpGs of the individuals of the test set were compared to those found by generating the data from the latent embeddings.

*MethylNet* was then configured for regression tasks and applied to derive sample age estimates in the Johansson data, using the reported chronological age as the ground truth. These results were compared to those derived from the Hannum and Horvath clocks using *cgageR*[31, 32, 53]. The Shapley framework was employed to quantify the importance of the CpGs in making predictions for age across 8 different age groups split by 10-year increments. The CpG importance was compared between the groups through hierarchical clustering to find similarities between the age groups. The one thousand most important CpGs from each group were extracted and overlapped with CpGs defined by the Hannum model to depict the concordance of important CpGs between *MethylNet* and the Hannum model.

For a second task, *MethylNet* was configured for multi-target regression to estimate cell-type proportions. First, *estimateCellCounts2*, using the 450K legacy IDOL optimized library [12], was used to deconvolve the cell-type proportions from each sample to develop our best proxy to ground truth outcomes for training the model. The *MethylNet* model was trained on the *estimateCellCounts2* estimates of cell-type proportions for six different immune cell-types. *MethylNet* was then compared to results derived from applying the 350 IDOL derived CpGs legacy library from FlowSorted.Blood.EPIC[54] using two different deconvolution methods Robust Partial Correlations (RPC) and Cibersort implemented in *EpiDISH*[55]. The importance of each CpG to each cell-type was then quantified through SHAP. These Shapley coefficients were compared using hierarchical clustering. A similar clustering profile would indicate these cell-types share similar driving CpGs, and recovery of the cell-lineage dendrogram would demonstrate concordance with known biology. The one thousand most important CpGs from each cell-type were extracted and overlapped with the IDOL CpGs to inspect if the two models picked up similar cell-type-specific CpGs. Additional details regarding SHAP can be found in the supplementary material.

In the next task, *MethylNet* was used to classify samples to cancer types. The data for the classification task are from 8891 TCGA-acquired samples, representing 32 different cancer types (Supplementary Figure 1 and Supplementary Tables 1,2), and preprocessed using *PyMethylProcess* to yield a 200k CpG beta matrix. The features with the highest mean absolute deviation across samples were selected to both limit the computational complexity, memory of model training and capture the highest variation in the data. The highly variable sites are assumed to be more biologically meaningful than the lower variable sites. The *MethylNet* analysis pipeline was conducted on the pan-cancer dataset. The results from *MethylNet* were compared to a popular omics classification approach, a uniform manifold approximation and projection (UMAP) embedding of the samples, followed by support vector machine (SVM) classification. UMAP is an effective way to reduce the dimensionality of the data as well as preserve meaningful local and global structure in the data [56, 57]. Both were performed using *PyMethylProcess’s general_machine_learning* module, which executed a hyperparameter grid search of the SVM model. Finally, the embeddings of the different cancer subtypes were compared by calculating of the average cosine distance between clusters in the test samples. These distances were clustered using hierarchical clustering to form larger superclasses of cancer that demonstrate a shared embedding profile.

A sensitivity analysis was conducted to understand how *MethylNet* scales with number of training samples and features. The TCGA cohort dataset was utilized and split into *MethylationArray*s of increasing number of features, scaled almost logarithmically for low number of features and then number of features were scaled linearly. This generated sixteen separate datasets. These datasets were trained in parallel with 100-job hyperparameter scans to yield final predictions. The sensitivity analysis on training set size split up the training set into 10% increments from 10% to 100%, and each of the 10 sets were trained using 150-job hyperparameter scans. The number of training epochs was reduced to 50 for each analysis to limit the computational compute time.

Finally, a 100-job hyperparameter scan was conducted to predict smoking status on the Liu data. Gradient-based SHAPley estimates were acquired using SHAP. The CpG SHAP score for the test set samples were subset by the CpGs significantly associated with smoking identified by Joehanes et. al. 2016. The average rank of the highest absolute SHAPly score for each CpGs across individuals were compared to the rank of CpGs most significantly associated with smoking reported by Joehanes et. al. 2016. Correlation of these rank orders was determined through Pearson’s correlation coefficient and a non-correlation statistical test was employed to find a p-value for the relationship.

## Code Availability Statement

*MethylNet* was built using Python 3.6 and utilizes the PyTorch framework to run its deep learning models on GPUs using CUDA, although CPUs are also suppored. The workflow is available as an easily installable command line tool and API via PyPI as *methylnet* and on Docker [58] as *joshualevy44/methylnet*. The Docker image contains a test pipeline that requires one line to run through the hyperparameter training and evaluation of all framework components and can run on your local personal computer in addition to high performance computing. Help documentation, example scripts, and the analysis pipeline are available in the *MethylNet* GitHub repository (https://github.com/Christensen-Lab-Dartmouth/MethylNet).

Tests of our pipeline’s functionality can be conducted on Code Ocean at: https://doi.org/10.24433/CO.6373790.v1.

## List of Abbreviations

450K: HumanMethylation450
850K: HumanMethylationEPIC
ANN: Artificial Neural Networks
CpG: Cytosine-Guanine Dinucleotides
CWL: Common Workflow Language
DNAm: DNA Methylation
EWAS: Epigenome-Wide Association Studies
GPUs: Graphics Processing Units
L-DMR: Leukocyte Differentially Methylated Regions
RPMM: Recursively Partitioned Mixture Models
SHAP: Shapley Additive Feature Explanations
SVM: Support Vector Machine
TCGA: The Cancer Genome Atlas
UMAP: Uniform Manifold Approximation and Projection
VAE: Variational Auto-encoders

## Declarations

### Ethics Approval and Consent to Participate

Not applicable.

### Consent for Publication

Not applicable.

### Availability of Data and Materials

Data used in this study was acquired from GEO accessions GSE87571, GSE42861, and from The Cancer Genome Atlas (TCGA). Test data is available in our GitHub repository and the data can be tested using Code Ocean at: https://doi.org/10.24433/CO.6373790.v1.

### Correspondence Statement

Correspondence and request for materials should be addressed to Joshua J. Levy.

### Authors’ Contributions

The conception and design of the study were contributed by JJL and BCC. Implementation, programming, data acquisition, and analyses were by JJL. JJL and BCC wrote the manuscript and all authors contributed to writing and editing of the manuscript. CLP performed the EpiDISH comparisons. AJT, YC, CLP contributed towards refining the analytic plan and direction. AJT, YC, CLP, and JJL tested the pipeline. LAS provided technical support to streamline and debug important aspects of the pipeline.

## Acknowledgements

Not applicable.

