## Supplementary Material for "MethylNet: An Automated and Modular Deep Learning Approach for DNA Methylation Analysis"

### Dataset Statistics and Demographics

Supplementary Table 1: Male to Female Ratio Across Both Datasets

| | Train | Val | Test |
|---|---|---|---|
| **Pan-Cancer** | 2530:2360 | 355:346 | 713:686 |
| **Age/Cell-Type** | 240:262 | 36:35 | 59:85 |

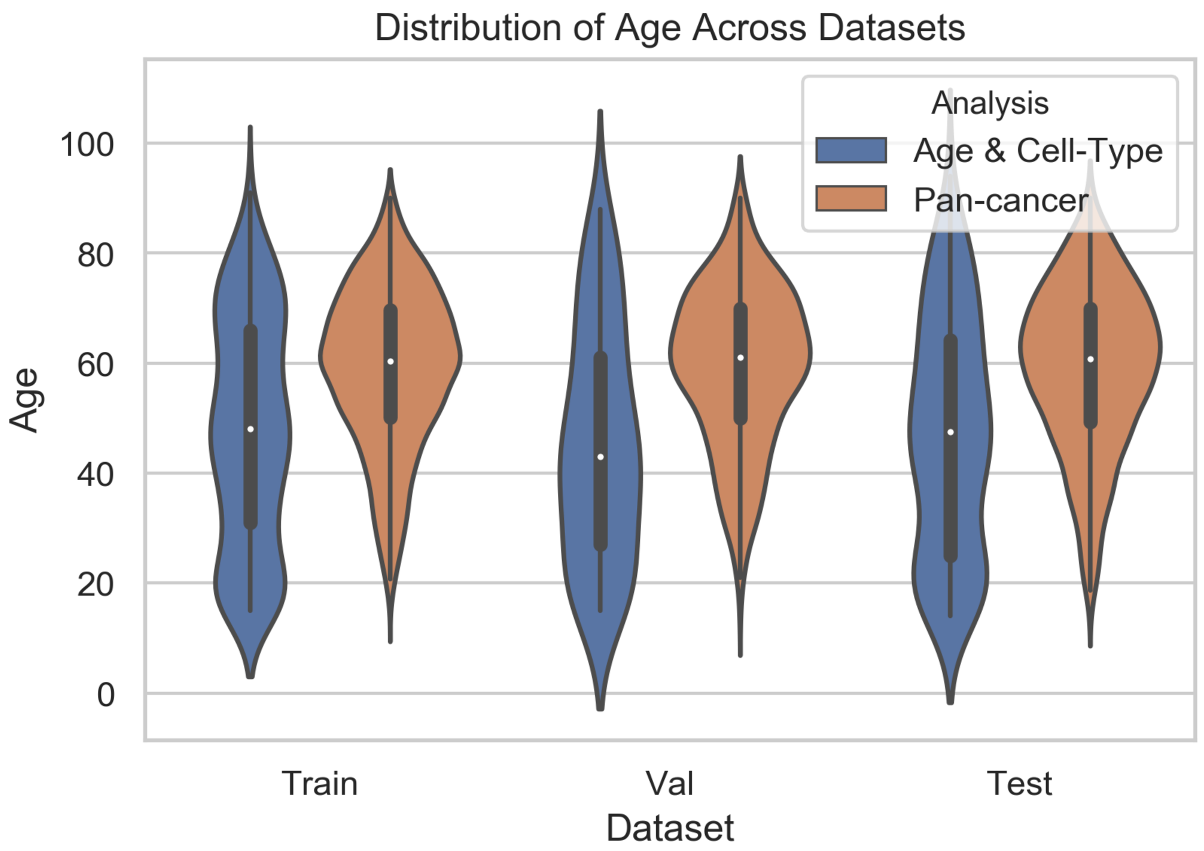

**Supplementary Figure 1: Distribution of Age Across Datasets**

Supplementary Table 2: Number of Samples for Each Cancer Subtype in Training, Validation, and Test Sets

| | Train | Val | Test |
|---|---|---|---|
| **BRCA** | 528 | 75 | 151 |
| **HNSC** | 355 | 51 | 101 |
| **LGG** | 347 | 50 | 99 |
| **THCA** | 346 | 50 | 99 |
| **PRAD** | 341 | 49 | 97 |
| **LUAD** | 311 | 44 | 89 |
| **SKCM** | 304 | 44 | 87 |
| **UCEC** | 298 | 42 | 85 |
| **BLCA** | 271 | 39 | 77 |
| **STAD** | 266 | 38 | 76 |
| **LIHC** | 247 | 35 | 70 |
| **LUSC** | 247 | 35 | 70 |
| **KIRC** | 216 | 31 | 62 |
| **CESC** | 206 | 30 | 59 |
| **COAD** | 180 | 26 | 52 |
| **SARC** | 167 | 24 | 48 |
| **KIRP** | 153 | 22 | 44 |
| **LAML** | 129 | 18 | 37 |
| **PCPG** | 122 | 18 | 35 |
| **ESCA** | 117 | 17 | 34 |
| **PAAD** | 110 | 16 | 32 |
| **GBM** | 96 | 14 | 27 |
| **TGCT** | 93 | 13 | 27 |
| **THYM** | 84 | 12 | 24 |
| **READ** | 62 | 9 | 18 |
| **MESO** | 61 | 9 | 17 |
| **ACC** | 49 | 7 | 14 |
| **UVM** | 44 | 6 | 13 |
| **KICH** | 37 | 5 | 11 |
| **DLBC** | 32 | 5 | 9 |
| **UCS** | 32 | 5 | 9 |
| **CHOL** | 9 | 1 | 3 |

### Unsupervised Analyses

a

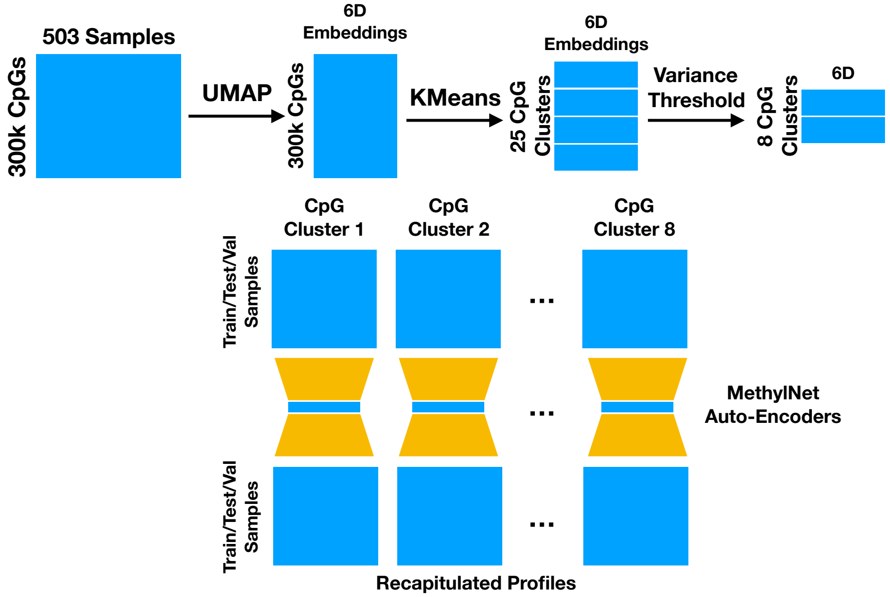

b

Recapitulation Scores for CpG Clusters on Held-Out Test Set
(n=144)

| CpG Cluster | Number CpGs | 6D-Cluster Variance | $R^2$ | Mean Absolute Residual |
|---|---|---|---|---|
| 23 | 15473 | 0.76 | 0.91 | 0.019 |
| 4 | 17184 | 0.76 | 0.93 | 0.017 |
| 1 | 13805 | 0.84 | 0.95 | 0.018 |
| 19 | 12835 | 0.88 | 0.92 | 0.018 |
| 11 | 12853 | 0.96 | 0.89 | 0.023 |
| 24 | 16381 | 0.99 | 0.95 | 0.02 |
| 12 | 5567 | 0.99 | 0.93 | 0.011 |
| 7 | 20018 | 1 | 0.91 | 0.023 |

**Supplementary Figure 2:** a) Visual flow diagram of method used to find CpG groupings and recapitulation of DNAm profiles. First, the 300k CpGs are projected into a 6-dimensional embedding using UMAP. Each point in the low dimensional space represents a CpG and proximity between points denotes a shared methylation profile across all of the training samples (n=503). Then, KMeans clustering was used to find 25 clusters of CpGs with similar profiles. The number of clusters of CpG features were reduced to 8 by filtering out clusters if their variance was above 1 in the 6D space. After that, the CpG features found in each cluster were used to select CpGs to form independent MethylationArrays across the training, validation and test sets. Finally, one autoencoder was trained per each array and the test samples were recapitulated and compared to the original input data; b) Descriptive statistics for final groupings of CpGs and recapitulation scores for each resultant set of CpGs versus the original methylation profiles input into each model.

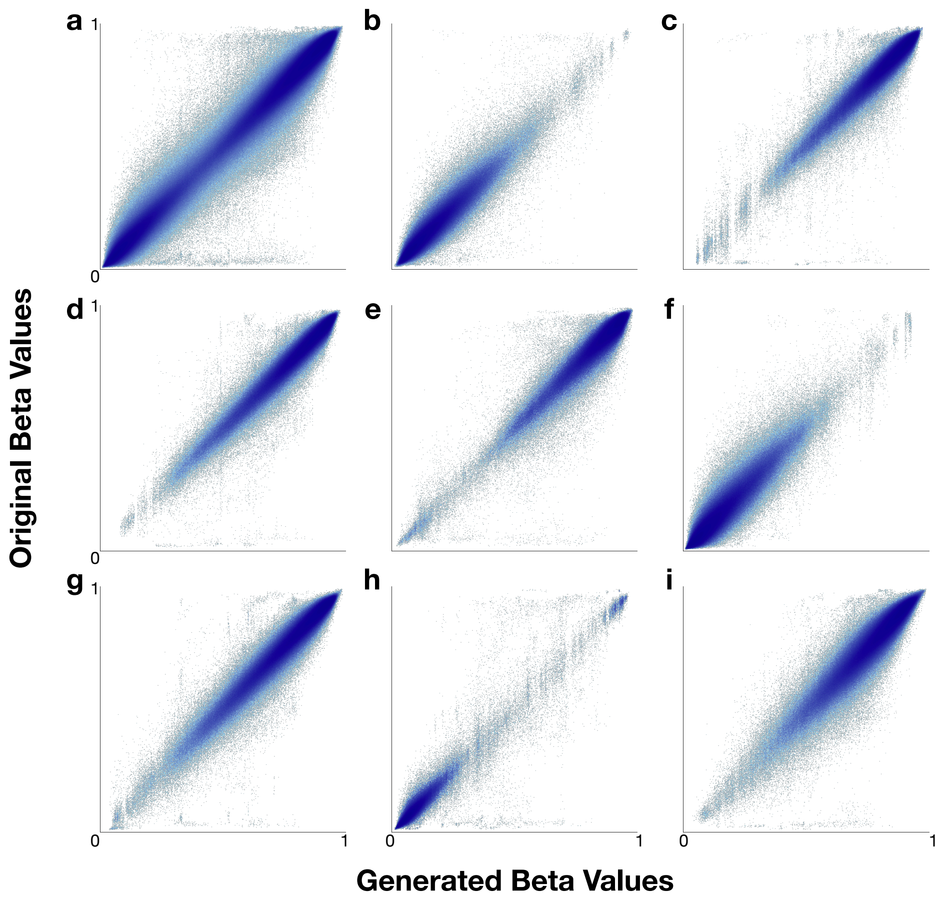

**Supplementary Figure 3:** Generated/recapitulated beta values versus original beta values for each CpG per individual of the held-out test set (n=144); b)-f) corresponds to each of eight chosen clusters in order of low to high cluster variance as previously described; a) is an aggregation of the generated/recapitulated versus true beta values of all of the CpG clusters

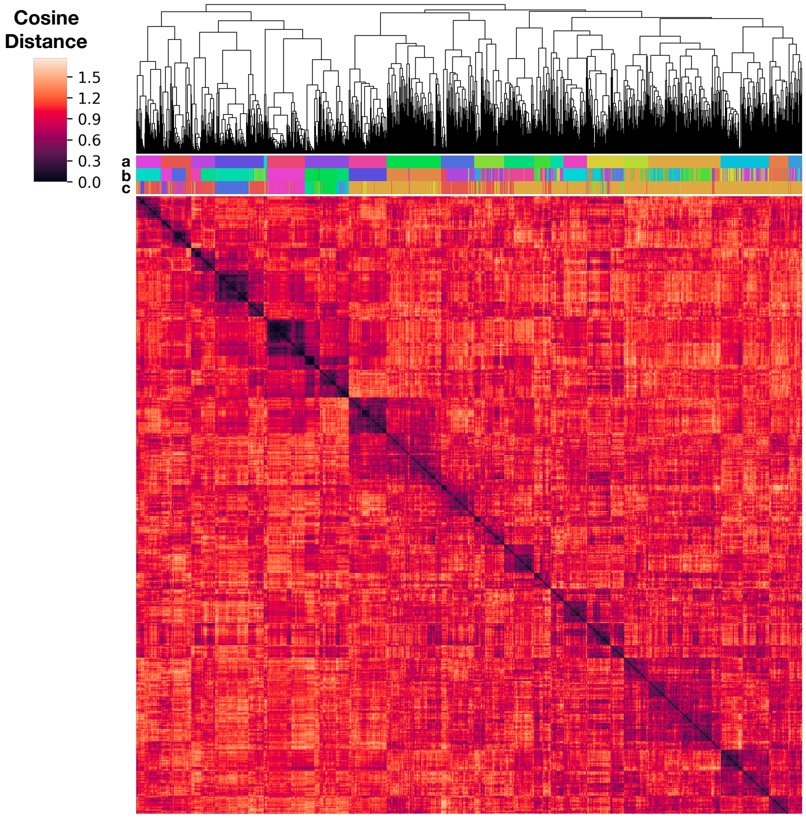

**Supplementary Figure 4:** Hierarchically clustered cosine distance matrix between test samples' VAE-embedded methylation profiles of the held-out test set for the TCGA cohort, colored by: a) Labels assigned to the hierarchical clustering labels for the samples; b) Original TCGA cancer labels; c) RPMM-derived clustering labels on 20k CpGs. Agreement scores between the RPMM and hierarchal clustering results and the original cancer subtypes were calculated using the v-measure, which takes into account the homogeneity and completeness of the labeling. Note that the clustering colors are not the same because the number of clusters is different from the number of cancer labels.

### Training and Validation Results

Supplementary Table 3: MethylNet Results on Training (n=503) and Validation (n=72) Sets for Sample Age Prediction

| Dataset | $R^2$ (mean±SE) | Mean Absolute Residual (mean±SE) | Explained Variance Proportion (mean±SE) |
|---|---|---|---|
| **Training** | 0.993±2.57e-04 | 1.49±0.0441 | 0.993±2.45e-04 |
| **Validation** | 0.973±0.00506 | 2.69±0.247 | 0.973±0.00505 |

Supplementary Table 4: MethylNet Results on Training (n=503) and Validation (n=72) Sets for Cell Type Deconvolution

| Cell Type | Dataset | $R^2$ (mean±SE) | Mean Absolute Residual (mean±SE) | Explained Variance Proportion (mean±SE) |
|---|---|---|---|---|
| **B Cell** | **Train** | 0.884±0.00967 | 0.00685±2.4e-04 | 0.89±0.00917 |
| | **Validation** | 0.756±0.0374 | 0.0101±9.88e-04 | 0.774±0.0371 |
| **CD4T** | **Train** | 0.975±0.00218 | 0.00628±1.79e-04 | 0.988±0.00102 |

| | | | | |
|---|---|---|---|---|
| | **Validation** | 0.874±0.0288 | 0.0121±0.00115 | 0.89±0.025 |
| **CD8T** | **Train** | 0.983±0.00152 | 0.00476±1.66e-04 | 0.986±0.00126 |
| | **Validation** | 0.758±0.0469 | 0.0154±0.00149 | 0.771±0.0388 |
| **Monocytes** | **Train** | 0.898±0.00947 | 0.00535±1.76e-04 | 0.898±0.00941 |
| | **Validation** | 0.385±0.0775 | 0.0122±0.00138 | 0.386±0.076 |
| **NK** | **Train** | 0.983±0.00151 | 0.0055±1.7e-04 | 0.99±9.69e-04 |
| | **Validation** | 0.819±0.0399 | 0.0121±0.00136 | 0.827±0.0373 |
| **Neutrophils** | **Train** | 0.995±4.44e-04 | 0.00524±1.67e-04 | 0.996±3.36e-04 |
| | **Validation** | 0.962±0.00942 | 0.012±0.0011 | 0.963±0.00851 |

Supplementary Table 5: MethylNet Results on Training (n=5860) and Validation (n=840) Sets for Pan-Cancer Classification

| Dataset | Accuracy Score | Recall Score | Precision Score | F1-Score |
|---|---|---|---|---|
| **Train** | 1.0±0 | 1.0±0 | 1.0±0 | 1.0±0 |
| **Validation** | 0.965±0.00622 | 0.965±0.00622 | 0.968±0.00524 | 0.966±0.00607 |

### SHAP Overlap with IDOL

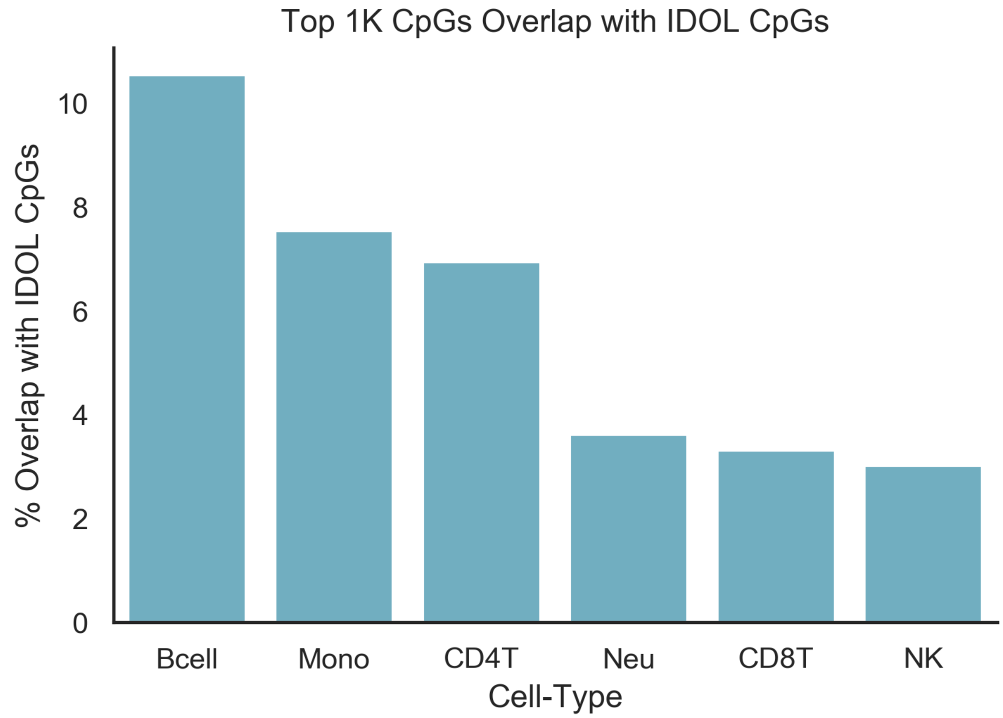

**Supplementary Figure 5:** Proportion of IDOL CpGs that are Overlapped by the Top 1k CpGs for Each Cell-Type

### Embedding Visualizations for Ages and Cell-Type Proportions

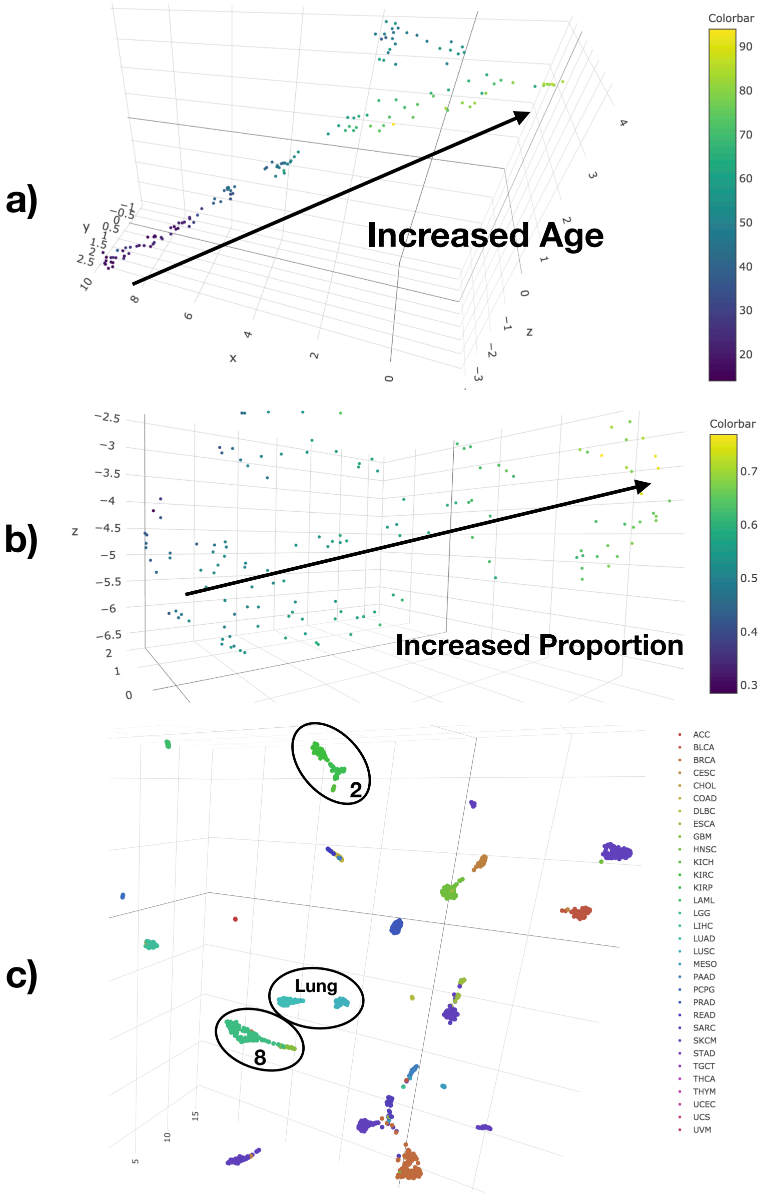

**Supplementary Figure 6:** Fine-tuned embeddings (few parts highlighted) for: a) Age Prediction, b) Cell-Type Deconvolution (colored by Neutrophil Cell Type Proportions), and c) Pan-Cancer Classification (Labeled kidney cancers, lung cancers, brain cancers)

Because of difficulties in interpretation associated with the distortion of perspective and UMAP parameters, these embeddings are also supplied as interactive three-dimensional plots and have been included in the GitHub repository at the following URL: https://github.com/Christensen-Lab-Dartmouth/MethylNet/tree/master/methylnet_results/embeddings

### Select Distributions of SHAPley Scores across Age and Cell-Type Groupings

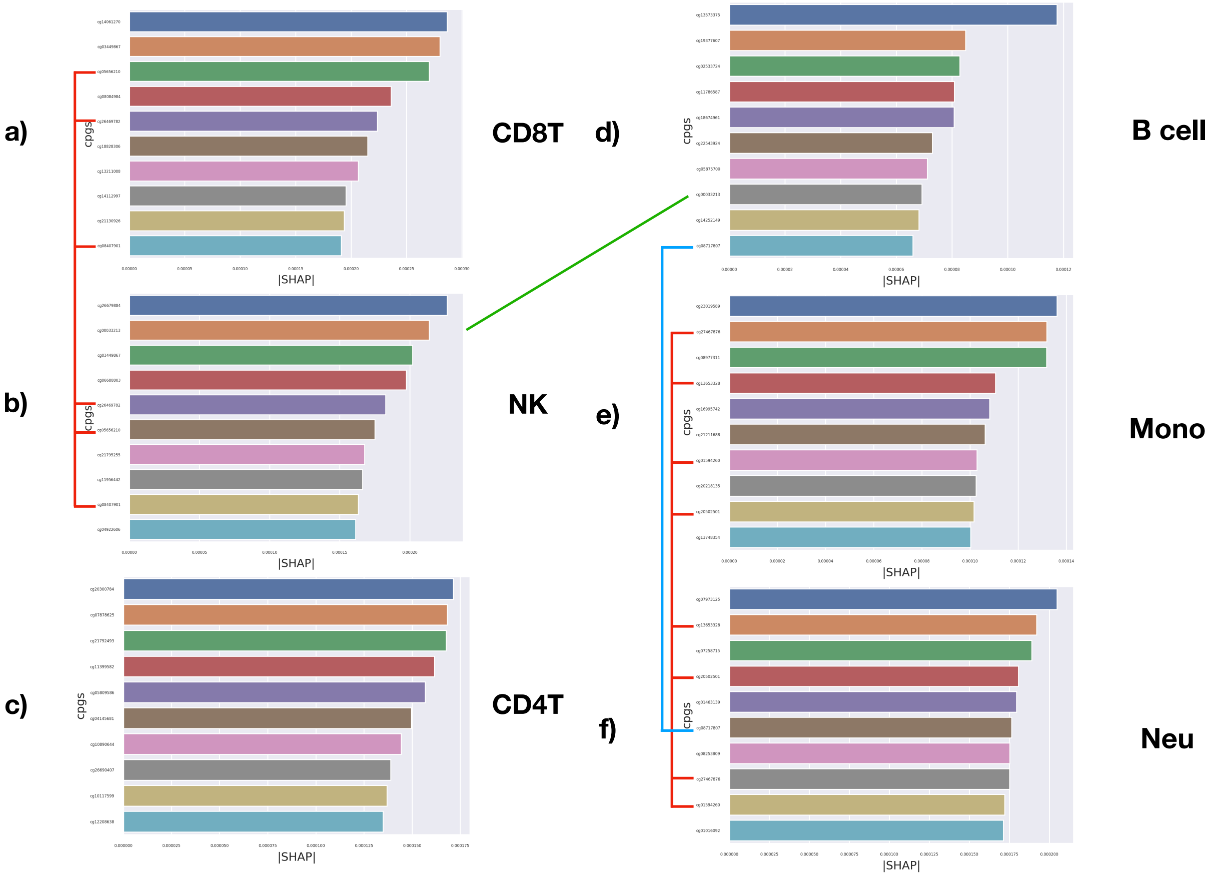

**Supplementary Figure 7:** Bar Charts of CpGs with the 10 Largest Shapley Scores for Each Cell-Type, linked by red, blue or green lines if shared across subtype for: a-d) Lymphocytes; e-f) Myeloids. Not sharing top 10 CpGs does not indicate that two cell-types do not share similar CpG profiles.

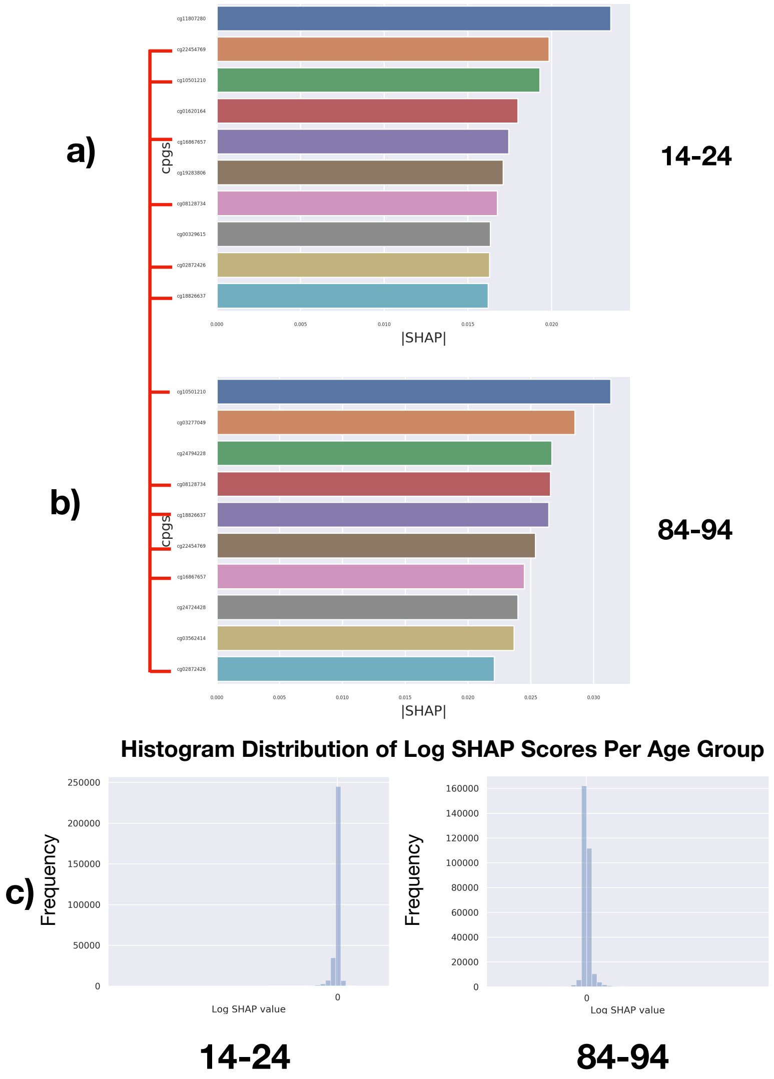

**Supplementary Figure 8:** Bar Charts of CpGs with the 10 Largest Shapley Scores for Age Groups: a) 14-24 and b) 84-94. These CpGs are linked if shared across the age groups, but this does not indicate that they are not shared outside of this top 10 list of CpGs. The top 10 CpGs that are associated with lower age are similar to the older age group; c) Distribution of Shapley Scores for these two age groups. CpG contributions tend to be negative for the younger age groups and positive for the older age groups.

### Confusion Matrix Pan-Cancer Classifications

Supplementary Table 6: Confusion Matrix Pan-cancer Classification (Colored Superclass)

| | | True Subtypes | | | | | | | | | | | | | | | | | | | | | | | | | | | | | | | |
|---|---|---|---|---|---|---|---|---|---|---|---|---|---|---|---|---|---|---|---|---|---|---|---|---|---|---|---|---|---|---|---|---|---|
| | | SARC | SKCM | UVM | BRCA | CHOL | LIHC | KICH | KIRC | KIRP | BLCA | CESC | UCEC | UCS | COAD | READ | LAML | ACC | PCPG | LUAD | MESO | THCA | ESCA | STAD | PAAD | PRAD | GBM | LGG | TGCT | THYM | HNSC | DLBC | LUSC |
| P | SARC | 45 | 0 | 0 | 0 | 0 | 0 | 0 | 0 | 0 | 1 | 0 | 0 | 0 | 0 | 0 | 0 | 0 | 0 | 0 | 2 | 0 | 0 | 0 | 0 | 0 | 1 | 0 | 0 | 0 | 0 | 0 | 0 |

| | | | | | | | | | | | | | | | | | | | | | | | | | | | | | | | | |
|---|---|---|---|---|---|---|---|---|---|---|---|---|---|---|---|---|---|---|---|---|---|---|---|---|---|---|---|---|---|---|---|---|
| SKCM | 1 | 87 | 0 | 1 | 0 | 0 | 0 | 0 | 0 | 0 | 0 | 0 | 0 | 0 | 0 | 0 | 0 | 0 | 0 | 0 | 0 | 0 | 0 | 0 | 0 | 0 | 0 | 0 | 0 | 0 | 0 | 0 |
| UVM | 0 | 0 | 13 | 0 | 0 | 0 | 0 | 0 | 0 | 0 | 0 | 0 | 0 | 0 | 0 | 0 | 0 | 0 | 0 | 0 | 0 | 0 | 0 | 0 | 0 | 0 | 0 | 0 | 0 | 0 | 0 | 0 |
| BRCA | 1 | 0 | 0 | 147 | 0 | 0 | 0 | 0 | 0 | 0 | 0 | 0 | 0 | 0 | 0 | 0 | 0 | 0 | 0 | 0 | 0 | 0 | 0 | 0 | 0 | 0 | 0 | 0 | 0 | 1 | 0 | 0 |
| CHOL | 0 | 0 | 0 | 0 | 3 | 1 | 0 | 0 | 0 | 0 | 0 | 0 | 0 | 0 | 0 | 0 | 0 | 0 | 0 | 0 | 0 | 0 | 0 | 0 | 0 | 0 | 0 | 0 | 0 | 0 | 0 | 0 |
| LIHC | 0 | 0 | 0 | 0 | 0 | 69 | 0 | 0 | 0 | 0 | 0 | 0 | 0 | 0 | 0 | 0 | 0 | 0 | 0 | 0 | 0 | 0 | 0 | 0 | 0 | 0 | 0 | 0 | 0 | 0 | 0 | 0 |
| KICH | 0 | 0 | 0 | 0 | 0 | 0 | 11 | 0 | 0 | 0 | 0 | 0 | 0 | 0 | 0 | 0 | 0 | 0 | 0 | 0 | 0 | 0 | 0 | 0 | 0 | 0 | 0 | 0 | 0 | 0 | 0 | 0 |
| KIRC | 0 | 0 | 0 | 0 | 0 | 0 | 0 | 60 | 2 | 0 | 0 | 0 | 0 | 0 | 0 | 0 | 0 | 0 | 0 | 0 | 0 | 0 | 0 | 0 | 0 | 0 | 0 | 0 | 0 | 0 | 0 | 0 |
| KIRP | 0 | 0 | 0 | 0 | 0 | 0 | 0 | 2 | 41 | 0 | 0 | 0 | 0 | 0 | 0 | 0 | 0 | 0 | 0 | 0 | 0 | 0 | 0 | 0 | 0 | 0 | 0 | 0 | 0 | 0 | 0 | 0 |
| BLCA | 0 | 0 | 0 | 0 | 0 | 0 | 0 | 0 | 1 | 75 | 0 | 0 | 0 | 0 | 0 | 0 | 0 | 0 | 0 | 0 | 0 | 0 | 0 | 0 | 0 | 0 | 0 | 0 | 0 | 0 | 0 | 0 |
| CESC | 0 | 0 | 0 | 0 | 0 | 0 | 0 | 0 | 0 | 1 | 57 | 0 | 0 | 0 | 0 | 0 | 0 | 0 | 0 | 0 | 0 | 0 | 0 | 0 | 0 | 0 | 0 | 0 | 0 | 1 | 0 | 0 |
| UCEC | 1 | 0 | 0 | 0 | 0 | 0 | 0 | 0 | 0 | 0 | 2 | 84 | 0 | 0 | 0 | 0 | 0 | 0 | 0 | 0 | 0 | 0 | 0 | 0 | 0 | 0 | 0 | 0 | 0 | 0 | 0 | 0 |
| UCS | 0 | 0 | 0 | 1 | 0 | 0 | 0 | 0 | 0 | 0 | 0 | 0 | 9 | 0 | 0 | 0 | 0 | 0 | 1 | 0 | 0 | 0 | 0 | 0 | 0 | 0 | 0 | 0 | 0 | 0 | 0 | 0 |
| COAD | 0 | 0 | 0 | 0 | 0 | 0 | 0 | 0 | 0 | 0 | 0 | 0 | 0 | 49 | 2 | 0 | 0 | 0 | 0 | 0 | 0 | 0 | 0 | 1 | 0 | 0 | 0 | 0 | 0 | 0 | 0 | 0 |
| READ | 0 | 0 | 0 | 0 | 0 | 0 | 0 | 0 | 0 | 0 | 0 | 0 | 0 | 3 | 16 | 0 | 0 | 0 | 0 | 0 | 0 | 0 | 0 | 1 | 0 | 0 | 0 | 0 | 0 | 0 | 0 | 0 |
| LAML | 0 | 0 | 0 | 0 | 0 | 0 | 0 | 0 | 0 | 0 | 0 | 0 | 0 | 0 | 0 | 37 | 0 | 0 | 0 | 0 | 0 | 0 | 0 | 0 | 0 | 0 | 0 | 0 | 0 | 0 | 0 | 0 |
| ACC | 0 | 0 | 0 | 0 | 0 | 0 | 0 | 0 | 0 | 0 | 0 | 0 | 0 | 0 | 0 | 0 | 14 | 0 | 0 | 0 | 0 | 0 | 0 | 0 | 0 | 0 | 0 | 0 | 0 | 0 | 0 | 0 |
| PCPG | 0 | 0 | 0 | 0 | 0 | 0 | 0 | 0 | 0 | 0 | 0 | 0 | 0 | 0 | 0 | 0 | 0 | 34 | 0 | 0 | 0 | 0 | 0 | 0 | 0 | 0 | 0 | 0 | 0 | 0 | 0 | 0 |
| LUAD | 0 | 0 | 0 | 0 | 0 | 0 | 0 | 0 | 0 | 0 | 0 | 0 | 0 | 0 | 0 | 0 | 0 | 0 | 87 | 0 | 0 | 0 | 0 | 0 | 0 | 0 | 0 | 0 | 0 | 0 | 0 | 4 |
| MESO | 0 | 0 | 0 | 1 | 0 | 0 | 0 | 0 | 0 | 0 | 0 | 1 | 0 | 0 | 0 | 0 | 0 | 0 | 0 | 15 | 0 | 0 | 0 | 1 | 0 | 0 | 0 | 0 | 0 | 0 | 0 | 0 |
| THCA | 0 | 0 | 0 | 0 | 0 | 0 | 0 | 0 | 0 | 0 | 0 | 0 | 0 | 0 | 0 | 0 | 0 | 0 | 0 | 0 | 99 | 0 | 0 | 0 | 0 | 0 | 0 | 0 | 0 | 1 | 0 | 0 |
| ESCA | 0 | 0 | 0 | 0 | 0 | 0 | 0 | 0 | 0 | 0 | 0 | 0 | 0 | 0 | 0 | 0 | 0 | 0 | 0 | 0 | 0 | 29 | 3 | 0 | 0 | 0 | 0 | 0 | 0 | 1 | 0 | 0 |
| STAD | 0 | 0 | 0 | 0 | 0 | 0 | 0 | 0 | 0 | 0 | 0 | 0 | 0 | 0 | 0 | 0 | 0 | 0 | 0 | 0 | 0 | 4 | 72 | 0 | 0 | 0 | 0 | 0 | 0 | 0 | 0 | 0 |
| PAAD | 0 | 0 | 0 | 0 | 0 | 0 | 0 | 0 | 0 | 0 | 0 | 0 | 0 | 0 | 0 | 0 | 0 | 1 | 0 | 0 | 0 | 0 | 0 | 29 | 0 | 0 | 0 | 0 | 0 | 0 | 0 | 0 |
| PRAD | 0 | 0 | 0 | 0 | 0 | 0 | 0 | 0 | 0 | 0 | 0 | 0 | 0 | 0 | 0 | 0 | 0 | 0 | 0 | 0 | 0 | 0 | 0 | 0 | 97 | 0 | 0 | 0 | 0 | 0 | 0 | 0 |
| GBM | 0 | 0 | 0 | 0 | 0 | 0 | 0 | 0 | 0 | 0 | 0 | 0 | 0 | 0 | 0 | 0 | 0 | 0 | 0 | 0 | 0 | 0 | 0 | 0 | 0 | 25 | 7 | 0 | 0 | 0 | 0 | 0 |
| LGG | 0 | 0 | 0 | 1 | 0 | 0 | 0 | 0 | 0 | 0 | 0 | 0 | 0 | 0 | 0 | 0 | 0 | 0 | 0 | 0 | 0 | 0 | 0 | 0 | 0 | 1 | 92 | 0 | 0 | 0 | 0 | 0 |
| TGCT | 0 | 0 | 0 | 0 | 0 | 0 | 0 | 0 | 0 | 0 | 0 | 0 | 0 | 0 | 0 | 0 | 0 | 0 | 0 | 0 | 0 | 0 | 0 | 0 | 0 | 0 | 0 | 27 | 0 | 0 | 0 | 0 |
| THYM | 0 | 0 | 0 | 0 | 0 | 0 | 0 | 0 | 0 | 0 | 0 | 0 | 0 | 0 | 0 | 0 | 0 | 0 | 0 | 0 | 0 | 0 | 0 | 0 | 0 | 0 | 0 | 0 | 24 | 0 | 0 | 0 |
| HNSC | 0 | 0 | 0 | 0 | 0 | 0 | 0 | 0 | 0 | 0 | 0 | 0 | 0 | 0 | 0 | 0 | 0 | 0 | 0 | 0 | 0 | 1 | 0 | 0 | 0 | 0 | 0 | 0 | 0 | 97 | 0 | 1 |
| DLBC | 0 | 0 | 0 | 0 | 0 | 0 | 0 | 0 | 0 | 0 | 0 | 0 | 0 | 0 | 0 | 0 | 0 | 0 | 0 | 0 | 0 | 0 | 0 | 0 | 0 | 0 | 0 | 0 | 0 | 0 | 9 | 0 |
| LUSC | 0 | 0 | 0 | 0 | 0 | 0 | 0 | 0 | 0 | 0 | 0 | 0 | 0 | 0 | 0 | 0 | 0 | 0 | 1 | 0 | 0 | 0 | 1 | 0 | 0 | 0 | 0 | 0 | 0 | 0 | 0 | 65 |

### Breakdown of Pan-cancer Classifications

**Supplementary Table 7: Breakdown Pancancer Classification Results (Colored by Superclass)**

| | Accuracy-Score | F1-Score |
|---|---|---|
| SARC | 0.94 | 0.97 |
| SKCM | 1.00 | 1.00 |
| UVM | 1.00 | 1.00 |
| BRCA | 0.97 | 0.99 |
| CHOL | 1.00 | 1.00 |
| LIHC | 0.99 | 0.99 |
| KICH | 1.00 | 1.00 |
| KIRC | 0.97 | 0.98 |
| KIRP | 0.93 | 0.96 |
| BLCA | 0.97 | 0.99 |
| CESC | 0.97 | 0.98 |
| UCEC | 0.99 | 0.99 |
| UCS | 1.00 | 1.00 |
| COAD | 0.94 | 0.97 |
| READ | 0.89 | 0.94 |
| LAML | 1.00 | 1.00 |
| ACC | 1.00 | 1.00 |
| PCPG | 0.97 | 0.99 |
| LUAD | 0.98 | 0.99 |
| MESO | 0.88 | 0.94 |
| THCA | 1.00 | 1.00 |
| ESCA | 0.85 | 0.92 |
| STAD | 0.95 | 0.97 |
| PAAD | 0.91 | 0.95 |
| PRAD | 1.00 | 1.00 |
| GBM | 0.93 | 0.96 |
| LGG | 0.93 | 0.96 |
| TGCT | 1.00 | 1.00 |
| THYM | 1.00 | 1.00 |
| HNSC | 0.96 | 0.98 |
| DLBC | 1.00 | 1.00 |
| LUSC | 0.93 | 0.96 |

### Average Cosine Distance Matrix between Cancer Subtypes for Pancancer-Embeddings

Supplementary Table 8: Average Cosine Distance Between Embeddings of Cancer Subtypes

| Subtypes | SARC | SKCM | UVM | BRCA | CHOL | LIHC | KICH | KIRC | KIRP | BLCA | CESC | UCEC | UCS | COAD | READ | LAML | ACC | PCPG | LUAD | MESO | THCA | ESCA | STAD | PAAD | PRAD | GBM | LGG | TGCT | THYM | HNSC | DLBC | LUSC |
|---|---|---|---|---|---|---|---|---|---|---|---|---|---|---|---|---|---|---|---|---|---|---|---|---|---|---|---|---|---|---|---|---|
| SARC | 0.0 | 0.7 | 0.7 | 1.0 | 0.8 | 1.0 | 0.7 | 0.9 | 1.0 | 1.0 | 0.8 | 0.8 | 0.7 | 1.2 | 0.9 | 0.9 | 0.8 | 0.9 | 1.4 | 0.6 | 0.9 | 0.9 | 0.8 | 0.7 | 1.2 | 0.8 | 0.9 | 0.9 | 0.9 | 0.9 | 0.7 | 1.1 |
| SKCM | 0.7 | 0.0 | 0.6 | 0.9 | 1.0 | 0.8 | 1.0 | 1.0 | 0.9 | 1.1 | 1.2 | 0.8 | 0.8 | 1.1 | 1.3 | 0.7 | 0.8 | 0.9 | 0.9 | 0.8 | 0.8 | 1.2 | 1.1 | 1.3 | 1.7 | 1.0 | 0.7 | 0.9 | 0.9 | 1.0 | 0.8 | 1.0 |
| UVM | 0.7 | 0.6 | 0.0 | 0.8 | 0.9 | 0.7 | 1.2 | 0.7 | 0.8 | 1.6 | 1.2 | 0.8 | 1.3 | 0.9 | 1.0 | 0.7 | 1.0 | 0.7 | 1.4 | 1.0 | 1.0 | 1.2 | 1.3 | 1.2 | 1.4 | 1.1 | 0.8 | 1.0 | 0.7 | 1.0 | 1.0 | 1.5 |
| BRCA | 1.0 | 0.9 | 0.8 | 0.0 | 0.8 | 0.9 | 1.3 | 1.5 | 1.2 | 1.3 | 0.9 | 0.9 | 0.9 | 0.9 | 1.1 | 1.1 | 0.9 | 0.9 | 0.9 | 0.8 | 1.3 | 0.9 | 1.1 | 1.1 | 1.1 | 1.4 | 1.0 | 1.0 | 0.8 | 1.2 | 1.4 | 1.3 |
| CHOL | 0.8 | 1.0 | 0.9 | 0.8 | 0.0 | 0.3 | 0.7 | 1.0 | 0.9 | 1.1 | 0.6 | 0.9 | 1.0 | 1.1 | 0.9 | 1.3 | 1.0 | 1.3 | 0.9 | 0.7 | 0.7 | 0.9 | 0.8 | 0.8 | 0.9 | 1.4 | 1.2 | 1.2 | 0.9 | 1.3 | 0.7 | 1.0 |
| LIHC | 1.0 | 0.8 | 0.7 | 0.9 | 0.3 | 0.0 | 0.8 | 0.7 | 0.9 | 1.1 | 1.0 | 0.8 | 1.2 | 1.0 | 0.9 | 1.1 | 0.9 | 0.9 | 0.8 | 1.0 | 0.9 | 1.2 | 0.9 | 0.9 | 1.2 | 1.4 | 1.3 | 0.8 | 0.8 | 1.2 | 0.7 | 1.0 |
| KICH | 0.7 | 1.0 | 1.2 | 1.3 | 0.7 | 0.8 | 0.0 | 0.6 | 0.6 | 0.7 | 1.1 | 1.0 | 0.7 | 1.3 | 0.9 | 1.3 | 0.7 | 1.0 | 0.9 | 0.8 | 0.7 | 0.9 | 0.7 | 0.7 | 0.7 | 0.7 | 0.8 | 1.1 | 1.3 | 1.3 | 0.8 | 0.8 |
| KIRC | 0.9 | 1.0 | 0.7 | 1.5 | 1.0 | 0.7 | 0.6 | 0.0 | 0.5 | 0.9 | 1.3 | 0.8 | 1.2 | 0.9 | 0.7 | 0.9 | 1.0 | 0.8 | 1.2 | 1.3 | 0.8 | 1.2 | 0.9 | 0.9 | 0.9 | 0.7 | 1.0 | 0.7 | 1.1 | 0.8 | 0.6 | 1.0 |
| KIRP | 1.0 | 0.9 | 0.8 | 1.2 | 0.9 | 0.9 | 0.6 | 0.5 | 0.0 | 1.0 | 1.1 | 1.0 | 0.8 | 0.7 | 0.8 | 1.0 | 1.1 | 0.8 | 0.8 | 1.0 | 0.7 | 0.9 | 1.0 | 1.2 | 0.9 | 1.1 | 1.0 | 1.3 | 1.4 | 1.1 | 1.2 | 1.1 |
| BLCA | 1.0 | 1.1 | 1.6 | 1.3 | 1.1 | 1.1 | 0.7 | 0.9 | 1.0 | 0.0 | 0.6 | 0.8 | 0.7 | 1.0 | 0.7 | 0.8 | 0.8 | 0.9 | 0.8 | 0.9 | 0.7 | 1.0 | 0.9 | 0.7 | 0.7 | 0.8 | 1.4 | 0.9 | 1.2 | 0.9 | 0.8 | 0.6 |
| CESC | 0.8 | 1.2 | 1.2 | 0.9 | 0.6 | 1.0 | 1.1 | 1.3 | 1.1 | 0.6 | 0.0 | 0.7 | 0.7 | 0.9 | 0.7 | 1.0 | 1.2 | 1.2 | 1.1 | 0.8 | 0.8 | 0.8 | 1.0 | 0.8 | 0.8 | 1.3 | 1.5 | 1.2 | 1.0 | 0.8 | 0.9 | 1.0 |
| UCEC | 0.8 | 0.8 | 0.8 | 0.9 | 0.9 | 0.8 | 1.0 | 0.8 | 1.0 | 0.8 | 0.7 | 0.0 | 0.6 | 0.9 | 0.8 | 1.1 | 0.9 | 0.9 | 1.4 | 1.4 | 0.9 | 1.3 | 1.1 | 1.3 | 0.9 | 1.0 | 1.2 | 0.7 | 1.2 | 0.8 | 0.7 | 1.3 |
| UCS | 0.7 | 0.8 | 1.3 | 0.9 | 1.0 | 1.2 | 0.7 | 1.2 | 0.8 | 0.7 | 0.7 | 0.6 | 0.0 | 1.0 | 1.0 | 1.2 | 0.8 | 1.1 | 1.0 | 0.9 | 0.9 | 0.8 | 1.0 | 1.2 | 0.9 | 1.1 | 1.0 | 1.3 | 1.6 | 1.0 | 1.3 | 1.1 |
| COAD | 1.2 | 1.1 | 0.9 | 0.9 | 1.1 | 1.0 | 1.3 | 0.9 | 0.7 | 1.0 | 0.9 | 0.9 | 1.0 | 0.0 | 0.3 | 0.7 | 1.0 | 0.8 | 1.0 | 1.3 | 1.5 | 0.9 | 0.8 | 1.1 | 1.0 | 1.2 | 1.3 | 1.3 | 1.4 | 1.0 | 1.3 | 1.4 |
| READ | 0.9 | 1.3 | 1.0 | 1.1 | 0.9 | 0.9 | 0.9 | 0.7 | 0.8 | 0.7 | 0.7 | 0.8 | 1.0 | 0.3 | 0.0 | 0.7 | 0.8 | 0.9 | 1.2 | 1.2 | 1.4 | 1.0 | 0.7 | 0.7 | 0.8 | 1.0 | 1.4 | 1.3 | 1.3 | 1.0 | 0.9 | 1.3 |
| LAML | 0.9 | 0.7 | 0.7 | 1.1 | 1.3 | 1.1 | 1.3 | 0.9 | 1.0 | 0.8 | 1.0 | 1.1 | 1.2 | 0.7 | 0.7 | 0.0 | 0.7 | 0.7 | 0.9 | 0.8 | 1.0 | 1.2 | 1.2 | 0.9 | 1.4 | 0.7 | 0.9 | 1.2 | 0.8 | 1.0 | 0.9 | 1.1 |
| ACC | 0.8 | 0.8 | 1.0 | 0.9 | 1.0 | 0.9 | 0.7 | 1.0 | 1.1 | 0.8 | 1.2 | 0.9 | 0.8 | 1.0 | 0.8 | 0.7 | 0.0 | 0.6 | 0.8 | 0.9 | 1.1 | 1.4 | 1.0 | 0.7 | 1.1 | 0.8 | 0.8 | 1.1 | 1.1 | 1.6 | 1.1 | 1.2 |
| PCPG | 0.9 | 0.9 | 0.7 | 0.9 | 1.3 | 0.9 | 1.0 | 0.8 | 0.8 | 0.9 | 1.2 | 0.9 | 1.1 | 0.8 | 0.9 | 0.7 | 0.6 | 0.0 | 1.0 | 0.8 | 0.9 | 1.3 | 1.4 | 0.9 | 1.0 | 1.0 | 1.1 | 0.8 | 0.8 | 1.2 | 1.3 | 1.3 |
| LUAD | 1.4 | 0.9 | 1.4 | 0.9 | 0.9 | 0.8 | 0.9 | 1.2 | 0.8 | 0.8 | 1.1 | 1.4 | 1.0 | 1.0 | 1.2 | 0.9 | 0.8 | 1.0 | 0.0 | 0.7 | 0.8 | 0.8 | 1.0 | 1.0 | 1.0 | 1.2 | 1.0 | 1.2 | 1.0 | 1.3 | 1.2 | **0.6** |
| MESO | 0.6 | 0.8 | 1.0 | 0.8 | 0.7 | 1.0 | 0.8 | 1.3 | 1.0 | 0.9 | 0.8 | 1.4 | 0.9 | 1.3 | 1.2 | 0.8 | 0.9 | 0.8 | 0.7 | 0.0 | 0.6 | 0.7 | 1.1 | 0.7 | 1.1 | 1.1 | 1.0 | 1.2 | 0.7 | 1.1 | 1.2 | 0.8 |
| THCA | 0.9 | 0.8 | 1.0 | 1.3 | 0.7 | 0.9 | 0.7 | 0.8 | 0.7 | 0.7 | 0.8 | 0.9 | 0.9 | 1.5 | 1.4 | 1.0 | 1.1 | 0.9 | 0.8 | 0.6 | 0.0 | 1.1 | 1.4 | 1.0 | 0.8 | 0.9 | 0.9 | 0.9 | 0.7 | 0.9 | 0.8 | 0.6 |
| ESCA | 0.9 | 1.2 | 1.2 | 0.9 | 0.9 | 1.2 | 0.9 | 1.2 | 0.9 | 1.0 | 0.8 | 1.3 | 0.8 | 0.9 | 1.0 | 1.2 | 1.4 | 1.3 | 0.8 | 0.7 | 1.1 | 0.0 | 0.5 | 0.8 | 0.8 | 1.1 | 1.0 | 1.1 | 1.1 | 0.7 | 1.2 | 0.7 |
| STAD | 0.8 | 1.1 | 1.3 | 1.1 | 0.8 | 0.9 | 0.7 | 0.9 | 1.0 | 0.9 | 1.0 | 1.1 | 1.0 | 0.8 | 0.7 | 1.2 | 1.0 | 1.4 | 1.0 | 1.1 | 1.4 | 0.5 | 0.0 | 0.6 | 0.9 | 0.8 | 0.9 | 0.9 | 1.2 | 0.8 | 0.7 | 0.7 |
| PAAD | 0.7 | 1.3 | 1.2 | 1.1 | 0.8 | 0.9 | 0.7 | 0.9 | 1.2 | 0.7 | 0.8 | 1.3 | 1.2 | 1.1 | 0.7 | 0.9 | 0.7 | 0.9 | 1.0 | 0.7 | 1.0 | 0.8 | 0.6 | 0.0 | 0.7 | 0.7 | 1.0 | 0.9 | 0.7 | 1.0 | 0.7 | 0.7 |
| PRAD | 1.2 | 1.7 | 1.4 | 1.1 | 0.9 | 1.2 | 0.7 | 0.9 | 0.9 | 0.7 | 0.8 | 0.9 | 0.9 | 1.0 | 0.8 | 1.4 | 1.1 | 1.0 | 1.0 | 1.1 | 0.8 | 0.8 | 0.9 | 0.7 | 0.0 | 0.9 | 1.0 | 1.0 | 1.1 | 1.0 | 1.1 | 0.9 |
| GBM | 0.8 | 1.0 | 1.1 | 1.4 | 1.4 | 1.4 | 0.7 | 0.7 | 1.1 | 0.8 | 1.3 | 1.0 | 1.1 | 1.2 | 1.0 | 0.7 | 0.8 | 1.0 | 1.2 | 1.1 | 0.9 | 1.1 | 0.8 | 0.7 | 0.9 | 0.0 | 0.4 | 0.8 | 0.9 | 0.7 | 0.6 | 0.7 |
| LGG | 0.9 | 0.7 | 0.8 | 1.0 | 1.2 | 1.3 | 0.8 | 1.0 | 1.0 | 1.4 | 1.5 | 1.2 | 1.0 | 1.3 | 1.4 | 0.9 | 0.8 | 1.1 | 1.0 | 1.0 | 0.9 | 1.0 | 0.9 | 1.0 | 1.0 | 0.4 | 0.0 | 1.0 | 0.9 | 1.0 | 0.9 | 0.8 |
| TGCT | 0.9 | 0.9 | 1.0 | 1.0 | 1.2 | 0.8 | 1.1 | 0.7 | 1.3 | 0.9 | 1.2 | 0.7 | 1.3 | 1.3 | 1.3 | 1.2 | 1.1 | 0.8 | 1.2 | 1.2 | 0.9 | 1.1 | 0.9 | 0.9 | 1.0 | 0.8 | 1.0 | 0.0 | 0.6 | 0.6 | 0.6 | 0.7 |
| THYM | 0.9 | 0.9 | 0.7 | 0.8 | 0.9 | 0.8 | 1.3 | 1.1 | 1.4 | 1.2 | 1.0 | 1.2 | 1.6 | 1.4 | 1.3 | 0.8 | 1.1 | 0.8 | 1.0 | 0.7 | 0.7 | 1.1 | 1.2 | 0.7 | 1.1 | 0.9 | 0.9 | 0.6 | 0.0 | 0.8 | 0.7 | 0.7 |
| HNSC | 0.9 | 1.0 | 1.0 | 1.2 | 1.3 | 1.2 | 1.3 | 0.8 | 1.1 | 0.9 | 0.8 | 0.8 | 1.0 | 1.0 | 1.0 | 1.0 | 1.6 | 1.2 | 1.3 | 1.1 | 0.9 | 0.7 | 0.8 | 1.0 | 1.0 | 0.7 | 1.0 | 0.6 | 0.8 | 0.0 | 0.7 | 0.7 |
| DLBC | 0.7 | 0.8 | 1.0 | 1.4 | 0.7 | 0.7 | 0.8 | 0.6 | 1.2 | 0.8 | 0.9 | 0.7 | 1.3 | 1.3 | 0.9 | 0.9 | 1.1 | 1.3 | 1.2 | 1.2 | 0.8 | 1.2 | 0.7 | 0.7 | 1.1 | 0.6 | 0.9 | 0.6 | 0.7 | 0.7 | 0.0 | 0.6 |
| LUSC | 1.1 | 1.0 | 1.5 | 1.3 | 1.0 | 1.0 | 0.8 | 1.0 | 1.1 | 0.6 | 1.0 | 1.3 | 1.1 | 1.4 | 1.3 | 1.1 | 1.2 | 1.3 | **0.6** | 0.8 | 0.6 | 0.7 | 0.7 | 0.7 | 0.9 | 0.7 | 0.8 | 0.7 | 0.7 | 0.7 | 0.6 | 0.0 |

### Dataset Scaling and Comparison with MLP

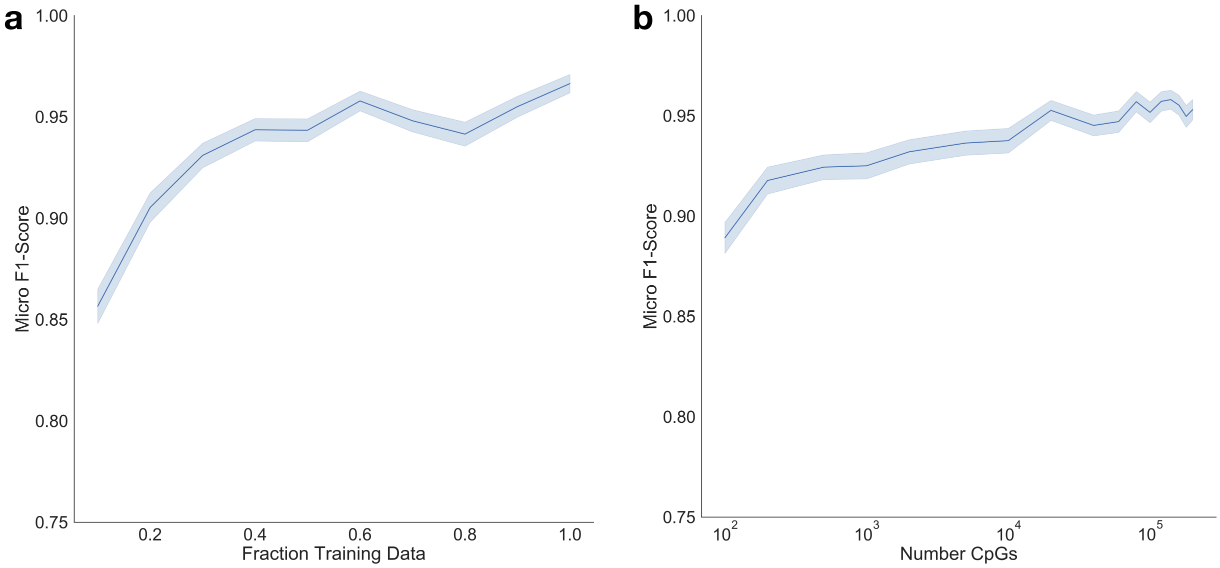

**c**

Comparison of Training MLP versus Pre-Training MethylNet with Auto-Encoder (n=1676)

| | F1-Score | First Epoch Val-Loss | Minimum Val-Loss |
|---|---|---|---|
| **Pre-trained** | 0.97±0.0045 | 19.03 | 2.37 |
| **MLP-Only** | 0.95±0.0051 | 33.75 | 3.33 |

**Supplementary Figure 9:** Micro F1-Scores of the held-out test samples (n=1676) of the TCGA cohort as they relate to: a) the fraction of training samples included for the training process, b) the number of CpGs. Test performance scales linearly with the number of training samples and logarithmically with the number of CpGs. Confidence intervals were calculated using a 1k nonparametric bootstrap of the test results for each dataset size point in the line plot, and the resulting bootstrapped f1-scores were used to compute the confidence interval for each point in the line plots; c) performance of MethylNet, pretrained using a VAE, is compared to performance using an MLP with the same architecture; F1-Score confidence intervals  were derived using a 1k nonparametric bootstrap; validation loss for each model is compared at the first training epoch and their ultimate convergence point.

### EWAS Analysis

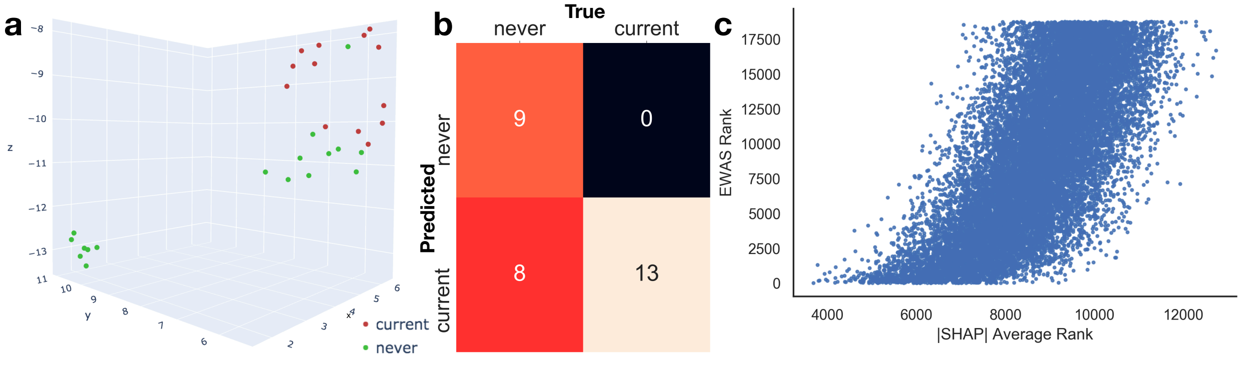

**Supplementary Figure 10:** Smoking EWAS study via MethylNet: a) final embeddings derived when finetuning the MethylNet VAE demonstrates cluster separation of the "never" versus "current" smokers; b) confusion matrix for the true and predicted "never" versus "current" smokers; c) plotted average ranks found using SHAP for the CpGs that intersected with CpGs identified by Liu et. al. versus the ranks of those corresponding p-values of the EWAS meta-analysis.

### Formulation of Variational Auto-Encoder

Variational Auto-encoders (VAE) were used to extract biologically meaningful features for downstream prediction tasks. The major components of these auto-encoders are the encoder and decoder neural networks. The encoder finds a low dimensional vector that is truthful to the original sample and the decoder up-samples this vector into a close approximation of the original sample. The encoder is represented by a function $q_\theta$ with neural network parameters $\theta$, and acts on input $x$ to produce the hidden representation $q_\theta(z|x)$. The decoder, $p_\phi$, with neural network parameters $\phi$, transforms $z$ into $\hat{x} = p_\phi(x|z)$, an approximation of $x$. The neural network trains off of the loss function: $l_{Recon}(\theta, \phi) = l_{Recon}\left(x, p_\phi\left(x|q_\theta(z|x)\right)\right)$. This loss function is known as the reconstruction loss and measures the difference between the original and decompressed sample. The loss function used to specify this difference is usually binary cross entropy loss or mean squared error.

The goal of an auto-encoder is to learn a compressed representation of the data by compressing using the encoder and decompressing using the decoder as aforementioned. Variational auto-encoders are generative models that seek to generate new data by sampling from some underlying distribution to make the distances between the compressed data meaningful. Therefore, in addition to learning to accurately reconstruct the original data, it also tries model the parameters of the encoder $q_\theta(z|x)$ to learn this probability distribution, $p(z)$, a multivariate distribution typically assumed to be gaussian, but is an active area of research to find a better prior. To do this, regularization is added to the loss function specified for the vanilla auto-encoder:

$$l(\theta, \phi) = l_{Recon}(\theta, \phi) + KL(q_\theta(z|x)||p(z))$$

Where $KL$ represents the KL-Divergence, which measures the difference between the encoder distribution $q_\theta(z|x)$ and the assumed latent distribution of data $p(z)$. This function penalizes any encoder representation that diverges from the specified probability distribution. To increase the generative qualities of the VAE, the $KL$ is usually given more weight, $\beta$, to more heavily penalize divergences from the distribution at the expense of sacrificing the ability to more accurately reconstruct the original samples. The following loss function was implemented in MethylNet:

$$l(\theta, \phi) = l_{Recon}(\theta, \phi) - \beta * l_{KL}(\theta)$$

Where $l_{KL}(\theta) = -KL(q_\theta(z|x)||p(z))$. $\beta$ was set to 1 for the training of the analyses mentioned in the paper in order make the validation losses of different hyperparameter runs comparable and to reduce the weight given to data generation, but this is a modifiable hyperparameter that can be altered during the optimization of the model. The encoder $q_\theta(z|x)$ was used for both the embedding training and fine-tuning steps of MethylNet's framework, but the decoder was discarded during prediction tasks. $p(z)$ was assumed to be multivariate standard normal in the MethylNet framework. A reparameterization trick samples this distribution during both the training of the feature extractor and training of the downstream prediction layers during finetuning, which serves the purpose of sampling a new $z$ at every training step. This augmentation technique may make the final neural network more generalizable to new data. The reparameterization was shutoff during test time.

### More Information on SHAPley Attribution Method

The SHAP (SHapley Additive ExPlanation) approach presents a possible framework to discover important CpGs for each prediction. They explain “black-box” models by simpler linear models. When applied to image analysis these methods translate the prediction of the model into heatmaps that overlay the original input image in a way that humans can understand. Applied to molecular profiles such as gene expression and CpG methylation information through interrogation of the coefficients of the simpler model, SHAP techniques can locate important genes and distinct CpGs that are drivers of association with an outcome variable.

The goal of Shapley feature attribution is to help the user understand why a machine learning model has made a specific prediction for any given sample. Many of these models are difficult to interpret. SHAP (Shapley Additive Explanations) is a model agnostic method to estimate the impact of each feature (assigned via Shapley values) on the prediction while remaining truthful to the properties of additivity, consistency and local accuracy.

Shapley values stem from game theory. A model’s output can be thought of as a reward to be distributed to a team of features that helped attain the reward. Shapley values dictate how the output of a model, the reward, should be shared amongst the features. Given the prediction of model $f$ on sample $x_i$ the method estimates shapley values $\phi(x_i, f)$ by finding an approximation to the model, $g_i(x_i)$, one model per sample $x_i$. This model is said to be locally accurate: its output converges to $f(x_i)$ for this sample by summing up (additivity) $J$ attributions $\phi_j(x_i, f)$ for $J$ features. It is also consistent: features that are truly important to one model’s predictions versus another are always assigned higher importance. To summarize these effects:

$$f(x_i) \approx g_i(x_i) = E[f(x_i)] + \sum_{j=1}^{J} \phi_j\,(x_i, f)$$

Where $E[f(x_i)]$ is the expected reward across the training samples, which was missing albeit nonessential from the above explanation. These Shapley values can be calculated by a weighted sum of the contribution of the feature to a model’s prediction given all possible permutations of other features being introduced into the model. This method is computationally intractable, so various estimation methods for deriving the Shapley values have been implemented. MethylNet employs kernel, gradient-based, and DeepLift SHAP inference methods to make quick approximations of these Shapley values.

*The SHAPley analysis was employed in the manuscript to study aging and cellular heterogeneity, and smoking. Given that $\phi_j$ can be calculated for each individual, we can produce a matrix of shapley values that can be summed over:*

$$\{\Phi_{ij} = \Phi_j(x_i, f)\}$$

*Then, we can calculate the Shapley values for each age group by summing the Shapley values across all of the individuals whose ages fall within the given ranges and arrive at aggregate measures for each age group k. A separate SHAP matrix is found for each cell-type, so an analogous task can be conducted by summing across all of the individuals for each cell type. Each age group was formed by binning the ages by discrete bin values (10 year increments), but Shapley values were calculated with respect to the prediction of a single individuals age and then averaged across the individuals of the group:*

$$\{\phi_{kj} = \left( \sum_{i \in \text{age group k}} \phi_j(x_i, f) \right) / \mathrm{N_k}\}$$

*Thus, the Shapley values were found by averaging attributions across* $\mathrm{N_k}$ *individuals of the same group, which for cell types is the entire set of individuals.*

*In Figure 2c in the text, we rank the Shapley values of each age group from maximum to minimum using order statistics:*

$$\phi_k^{(j)}$$

*Where (j) here is the jth largest Shapley value corresponding its respective feature and then look for the number of Hannum CpGs that overlap with the set of the top 1000* $\phi_k^{(j)}$*.*

*As for Figure 2d, we calculate the correlation distance between the Shapley values between pairs of age groups (i and jth age groups) as such:*

$$\mathrm{d_{ij}} = \mathrm{d}\left(\boldsymbol{\phi}_{k_i}, \boldsymbol{\phi}_{k_j}\right) = 1 - \frac{\left(\boldsymbol{\phi}_{k_i} - \overline{\boldsymbol{\phi}_{k_i}}\right) \cdot \left(\boldsymbol{\phi}_{k_j} - \overline{\boldsymbol{\phi}_{k_j}}\right)}{\left\|\left(\boldsymbol{\phi}_{k_i} - \overline{\boldsymbol{\phi}_{k_i}}\right)\right\|_2 \left\|\left(\boldsymbol{\phi}_{k_j} - \overline{\boldsymbol{\phi}_{k_j}}\right)\right\|_2}$$

*Where* $\boldsymbol{\phi}_{k_i}$ *is the sequence of shapley scores in the ith age group. Since due to high collinearity, it can be difficult to pinpoint exactly which CpGs are important in each age group, the correlation between the two age groups suggests that the two age groups have a similar overall profile of important CpGs that are associated with their outcome. Thus, clustering of these distances can yield relationships between the age groups that can corroborate with what we would expect in reality, and a similar analysis was conducted for cell-types in Figure 3c.*

*For the smoking comparison, we selected CpGs in our SHAP array that corresponded to the CpGs used in the cigarette smoking study, and then rank ordered the absolute value of each of the individual's CpG SHAP scores. We averaged the ranks of the CpGs across the individuals to ascertain the overall importance of the CpGs to compare to the ordering found from the traditional EWAS analysis.*

### Hyperparameter Scan Details for Embedding and Prediction Tasks

A robust hyperparameter scan was conducted for both the embedding and prediction tasks. A list of a user supplied search grid could be supplied; MethylNet also has its own built in search options. Typically, a number of randomized searches are made, the lowest validation losses from each of these runs are returned. Then, the top job(s) can be rerun to try to find the set of parameters with the lowest loss. Rerunning the best performing jobs may not yield the lowest validation loss due to the stochastic nature of initializing the neural network parameters and the samples shuffled during the training iterations, so it is recommended to rerun the analyses with the same set of hyperparameters multiple times. The following are a list of select hyperparameters that could be modified for these tasks:

- Number of epochs to train for
- Number of Latent Dimensions (Embedding Tasks only)
- Learning Rate (during prediction tasks, one is specified for feature extractor, the other for the fine-tuning layers)
- Beta (Embedding Tasks, weight given to KL-Loss)
- KL warm up (Embedding Task, number of epochs to apply lower weight to KL Loss to focus on reducing reconstruction loss)
- Scheduler (function that modulates the learning rate; for instance, interrupted cosine or horizontal line)
- Parameters relating to scaling of interrupted cosine learning rate scheduler
- Batch size
- Model Complexity (coefficient that determines how wide and deep neural network should be) and hidden layer topology (the topology of the encoder is mirrored to form the decoder)

Below are the top 3 unique sets of select hyperparameters for MethylNet's embedding and prediction tasks selected for the age estimation, cell-type deconvolution and pan-cancer classification analyses:

Supplementary Table 9: Select Hyperparameters for Embedding Tasks

| Task | Number Epochs | Best Epoch | Minimum Val Loss | Number Latent Dimensions | Encoder Hidden Layer Topology | Learning Rate | Batch Size |
|---|---|---|---|---|---|---|---|
| **Age Estimation** | 500 | 306 | 9159250 | 150 | [100, 200, 300] | 0.01 | 50 |
| | 200 | 40 | 9181492 | 100 | [500, 100] | 0.05 | 50 |
| | 200 | 193 | 9196771 | 100 | [500, 100] | 0.01 | 50 |
| **Cell-Type Deconvolution** | 500 | 246 | 9157337 | 100 | [300] | 0.005 | 50 |
| | 100 | 98 | 9160540 | 100 | [] | 0.01 | 50 |
| | 700 | 499 | 9169060 | 300 | [200, 1000, 100] | 0.1 | 50 |
| **Pan-Cancer** | 700 | 510 | 88319693 | 150 | [200, 100, 200] | 0.001 | 512 |
| | 500 | 305 | 88436906 | 100 | [500, 100] | 0.005 | 512 |
| | 700 | 636 | 88526436 | 100 | [300, 200] | 0.0005 | 256 |

Supplementary Table 10: Select Hyperparameters for Prediction Tasks

| Task | Number Epochs | Best Epoch | Minimum Validation Loss | Fine-Tuning Layer Topology | Learning Rate Feature Extractor | Learning Rate Fine-tuning Layers | Batch Size | Dropout |
|---|---|---|---|---|---|---|---|---|
| **Age Estimation** | 700 | 482 | 828.46 | [200] | 0.1 | 0.5 | 256 | 0 |
| | 200 | 173 | 942.90 | [] | 0.1 | 0.0001 | 100 | 0.2 |
| | 200 | 148 | 1029.74 | [100, 100, 200] | 0.1 | 0.0001 | 100 | 0 |
| **Cell-Type Deconvolution** | 700 | 630 | 0.12 | [1000] | 0.05 | 5E-05 | 256 | 0.1 |
| | 500 | 485 | 0.18 | [200, 500] | 0.1 | 5E-05 | 50 | 0 |
| | 500 | 479 | 0.24 | [500, 1000] | 0.0005 | 0.005 | 500 | 0 |
| **Pan-Cancer** | 200 | 126 | 2.19 | [300] | 0.001 | 1E-05 | 50 | 0 |
| | 500 | 256 | 2.22 | [] | 5E-05 | 0.0005 | 50 | 0 |
| | 500 | 204 | 2.31 | [200] | 0.001 | 5E-05 | 50 | 0.2 |

More information on the hyperparameter scans can be found in the supplied GitHub repository.

### Hyperparameter Scan Details for SVM Tasks

To compare MethylNet's Pan-Cancer predictions to another leading machine learning technique, a robust hyperparameter scan was conducted to find the best set of hyperparameters for an L2-penalized support vector machine model. Similar to MethylNet, the set of hyperparameters with the highest validation f1-score was chosen; class weights were used to account for any class imbalances. Listed below are the model's hyperparameters and their range:

- Kernel: linear or radial basis (RBF) decision functions
- Inverse weight given to L2-penalty (high value has low regularization): 1, 10, 100, 1000
- Gamma (influence of training sample for RBF kernel): 1, 0.1, 0.001, 0.0001

A randomized grid search that tested 24 sets of hyperparameters was conducted to find the best set of hyperparameters. This model was trained and tested using this set to produce the results from the paper.

### Example Code to Run Pipeline

Example code for running the pipeline can be found at https://github.com/Christensen-Lab-Dartmouth/MethylNet/tree/master/example_scripts. Further instructions can be found in the README and wiki page. The code can also be run on Code Ocean without install at: https://doi.org/10.24433/CO.6373790.v1 .

### Model Training Information and Training Curves

Binary Cross Entropy Loss was used as the loss function for the regression analyses, and Cross Entropy Loss was used for the classification approach. At every hidden layer in the network, non-linear transforms were applied via the ReLU transform, and during training, some of the neurons of these network layers were deactivated via Dropout for improved generalizability. The network was specified to use a sigmoid function to restrict the output of the regression tasks between 0 and 1. Throughout the training iterations, the learning rate was modulated using an interrupted cosine function to best capture trade-offs between exploration and exploitation of the objective. This functionality was sometimes removed at various points during the hyperparameter scans. The model with the lowest validation loss at a particular training iteration was returned for each run. The models were trained using a Nvidia Tesla K80 Graphics Processing Units, accessed via the Dartmouth Discovery Research Computing compute cluster.

Plotted below are the training curves of the auto-encoding and fine-tuning/transfer learning steps:

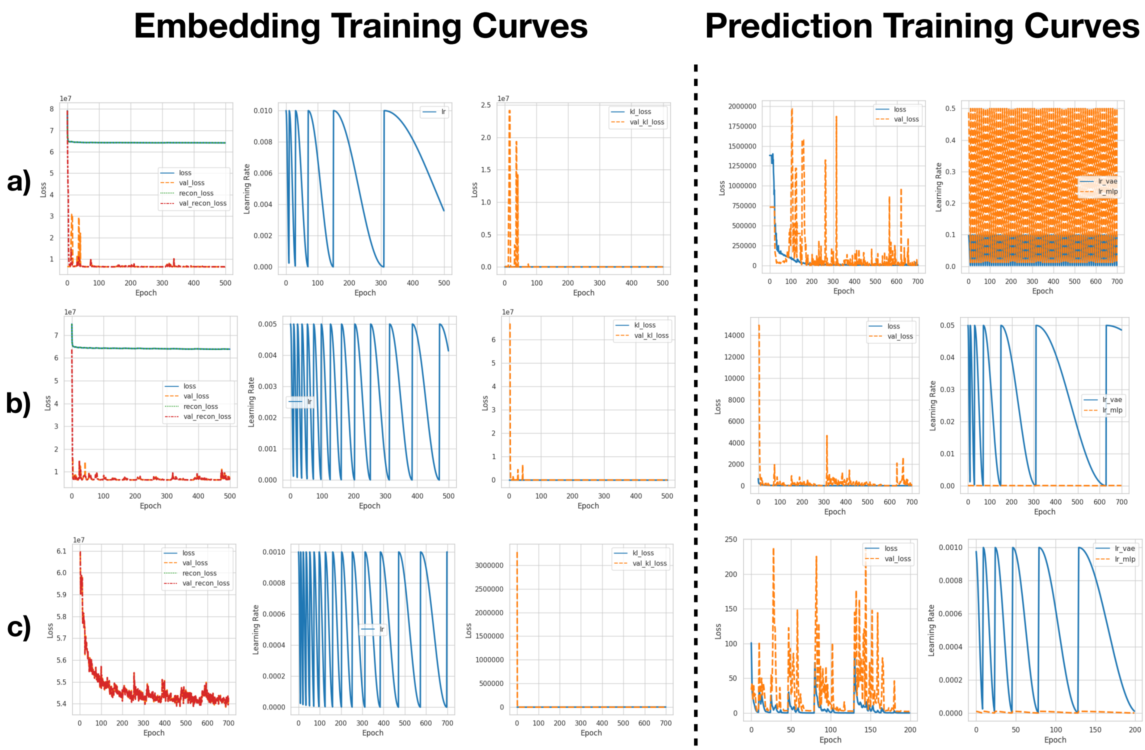

**Supplementary Figure 11: Model Training Curves for** a) Age Predictions, b) Cell-Type Predictions, c) Pan-Cancer Predictions. Please note that the learning rates for the prediction curve of a) oscillates quickly every 10 training epochs as compared to a larger timescale.
